# Classic oncogene family Myc defines unappreciated distinct lineage states of small cell lung cancer

**DOI:** 10.1101/852939

**Authors:** Ayushi S. Patel, Seungyeul Yoo, Ranran Kong, Takashi Sato, Abhilasha Sinha, Li Bao, Maya Fridrikh, Katsura Emoto, German Nudelman, Charles A. Powell, Mary Beth Beasley, Jun Zhu, Hideo Watanabe

## Abstract

Comprehensive genomic analyses of small cell lung cancer (SCLC), the most aggressive form of lung cancer, have revealed near universal loss of tumor suppressors *(RB1* and *TP53*) and frequent genomic amplification of all three *MYC* family members. The amplification of each Myc family member is mutually exclusive; hence it had been long suggested that they are functionally equivalent. However, their expression has more recently been associated with specific neuroendocrine markers and distinct histopathology. In this study, we explored a novel role of c-Myc and L-Myc as lineage determining factors contributing to SCLC molecular subtypes and histology. Integrated analyses of a gene regulatory network generated from mRNA expression of primary SCLC tumor and chromatin state profiling of SCLC cell lines showed that Myc family members impart distinct transcriptional programs associated with lineage state; wherein the L-Myc signature was enriched for neuronal pathways while the c-Myc signature was enriched for Notch signaling and epithelial-to-mesenchymal transition. We investigated the functional redundancy and distinction of c-Myc and L-Myc, and noted the insufficiency of L-Myc to induce lineage switch in contrast to the potential of c-Myc to induce trans-differentiation. c-Myc rewires the Myc-accessible landscape and activates neuron al repressor, Rest to mediate transition from ASCL1-SCLC to NeuroD1-SCLC characterized by distinct LCNEC-like histopathology. Collectively, our findings reveal a previously undescribed role of historically defined general oncogenes, c-Myc and L-Myc, for regulating lineage plasticity across molecular subtypes as well as histological subclasses.

## Introduction

Small cell lung cancer (SCLC) represents about 15% of all lung cancers with a median survival time of approximately 10 months and 5-year overall survival at 6% (*1*). SCLCs are characterized by neuroendocrine differentiation, rapid growth, early metastatic spread and poor prognosis (*2*). The standard of care has remained cytotoxic chemotherapy for decades, mainly etoposide combined with a platinum agent (*3*), that are only temporarily effective for the vast majority of patients (*4*). Even with recent developments in immune checkpoint inhibitors, activity against SCLCs when combined with chemotherapy has been marginal with a modest improvement in the median survival (10.3 months vs. 12.3 months) for extensive stage SCLCs treated with immunotherapy (*5*). These data reflect the urgent need for more effective therapeutics for patients with SCLC. The lack of effective therapeutics for SCLC stands in stark contrast to the breadth of targeted therapies for non-small cell lung cancer (NSCLC), particularly lung adenocarcinoma (*6*). The progress in drug development for NSCLCs is largely attributable to more comprehensive understanding of molecular subtypes and to identification of targetable driver oncogenes (*7*). Therefore, better characterization of the molecular subtypes of SCLC should aid future drug development and permit patient stratification for targeted therapies, a strategy that has been remarkably effective for specific subsets of advanced lung adenocarcinoma patients.

Characterization of SCLC subtypes was first noted by morphological differences over three decades ago when human SCLC cell lines were implanted as xenografts and distinguished as two primary subtypes: classical SCLC and variant SCLC (*8,9*). The classical subtype featured relatively small cells with high nuclear:cytoplasm ratio while the variant subtype exhibited relatively larger cells and moderate amounts of cytoplasm. However, the World Health Organization (WHO) classification, updated in 2015, histologically recognizes SCLC as a homogenous disease with neuroendocrine features defined by small cells, scant cytoplasm, nuclear morphology with fine granular chromatin and lacking prominent nucleoli, reminiscent of the features of the ‘classical’ SCLC. The originally described ‘variant’ subtype may represent combined SCLC with LCNEC in the current classification (*10*).

More recent efforts to distinguish SCLC molecular subtypes include profiling gene expression and genome-wide methylation in primary human tumors and patient derived xenografts (PDX). These profiles revealed three clusters, with a dichotomy between achaete-scute homologue 1 (ASCL1) and neurogenic differentiation factor 1 (NeuroD1) expression, in addition to a cluster with low expression of both (*11*). The expression of ASCL1 and NeuroD1 has been implicated to confer SCLC heterogeneity by imparting distinct transcriptional profiles (Borromeo et al., 2016). The third neuroendocrine-low cluster led to further classification into two subtypes characterized by transcriptional driver YAP1 or POU class 2 homeobox 3 (POU2F3) (*12,13*) SCLC cell line xenograft histology has been correlated with these contrasting factors, where variant SCLC was positively correlated with a higher NeuroD1:ASCL1 ratio and classical SCLC was positively correlated with a higher ASCL1:NeuroD1 ratio in SCLC cell lines (*11*).

Observations on Myc family characteristics in SCLC genetically engineered mouse models (GEMMs) has provided insight in their contribution to histopathological characteristics (*14,15*) The classical SCLC GEMM with conditional loss of *Rb1* and *Trp53* mouse, harbored stochastic *MYCL* amplifications or overexpression associated with classical SCLC histopathology (*16*). On the contrary, consistent with the original report on the variant subtype to harbor frequent *MYC* amplification (*9*), a more recent study showed that additional c-Myc overexpression, in the classical SCLC GEMM drives the progression of murine SCLC with variant histopathology and reduced neuroendocrine gene expression including ASCL1 but higher NeuroD1 expression (*17*). Yet, the molecular mechanisms underlying the distinction between L-Myc and c-Myc driven subsets of SCLC remain unexplored.

c-Myc (*MYC*) and L-Myc (*MYCL*) belong to the MYC family of basic helix-loop-helix (bHLH) transcription factors. The paralogs contain functionally relevant highly conserved amino acid sequences and are structurally homologous (*18,19*). c-Myc is a well-characterized oncogene; L-Myc although understudied is implicated to have a similar oncogenic role. Amplification of Myc family members is mutually exclusive and overall accounts for ~20% of SCLC and overexpression for ~50% of SCLC in primary human tumors (*20*). In contrast to the fact that *MYC* is commonly amplified across all three major lung cancer subtypes: lung adenocarcinomas, squamous cell lung carcinomas and SCLC (*20–22*), *MYCL* and *MYCN* are uniquely amplified in SCLC, in a manner suggestive of their role as lineage-amplified genes.

In this study, we explore a novel role of c-Myc and L-Myc as lineage specific factors to associate SCLC molecular subtypes with histological classes. We investigated the potential of L-Myc and c-Myc to regulate lineage state and identified transcriptional programs unique to each Myc family member, wherein L-Myc regulates neuronal developmental pathways and c-Myc regulates epithelial-to-mesenchymal transition and Notch signaling, biological pathways that are associated with distinct molecular subsets. In addition, we showed the requirement of c-Myc to maintain lineage state marker NeuroD1, in NeuroD1-positive SCLC and the incompatibility of c-Myc with ASCL1-positive SCLC that ultimately leads to trans-differentiation to NeuroD1-driven SCLC characterized by variant histopathology mediated by a transcriptional repressor, Rest.

## Results

### SCLC network reveals unique and distinct sub-networks for c-Myc and L-Myc

To understand biological processes that may be unique to c-Myc and L-Myc in SCLC, we sought to investigate potential causal regulations among genes. To this end, we first built a molecular causal network using primary SCLC datasets and focused on the networks that involve c-Myc and L-Myc. Combining two independent primary SCLC transcriptomic datasets (*20,23*), a Bayesian network was built using the software package RIMBANET we had previously developed (see Methods). The Bayesian network comprised of 8,451 unique genes (nodes) and 9,301 regulations (edges) among these genes. The regulations inferred in this Bayesian network provide insights into biological functions associated with molecular features such as pathways or gene signatures (*24,25*). This enables us to discern transcriptional sub-networks associated with c-Myc and L-Myc that may reflect their unique biological roles. We first aimed to generate a signature associated with each factor and then projected the signature to the SCLC Bayesian network to identify specific subnetworks (Figure 1A). To generate these signatures, we sought to select an independent dataset. Hence, we examined copy number alteration and expression for each Myc family member in 49 SCLC cell lines from Cancer Cell Line Encyclopedia (CCLE) and classified them into four groups representing each Myc family member and low Myc (Figure S1A). Then, we selected cell lines that belong to *MYC* and *MYCL* groups and examined mRNA expression *MYC* and *MYCL* to select cell lines for c-Myc with high expression of *MYC* and low expression of *MYCL* and vice versa (Figure S1B). We identified 457 differentially expressed genes (t-test p-value < 0.01 and fold change > 1.5); 147 and 310 genes overexpressed in *MYC* and *MYCL* SCLC cell lines respectively and defined them as their introductory gene signatures (Figure S1C).

**Figure 1:**
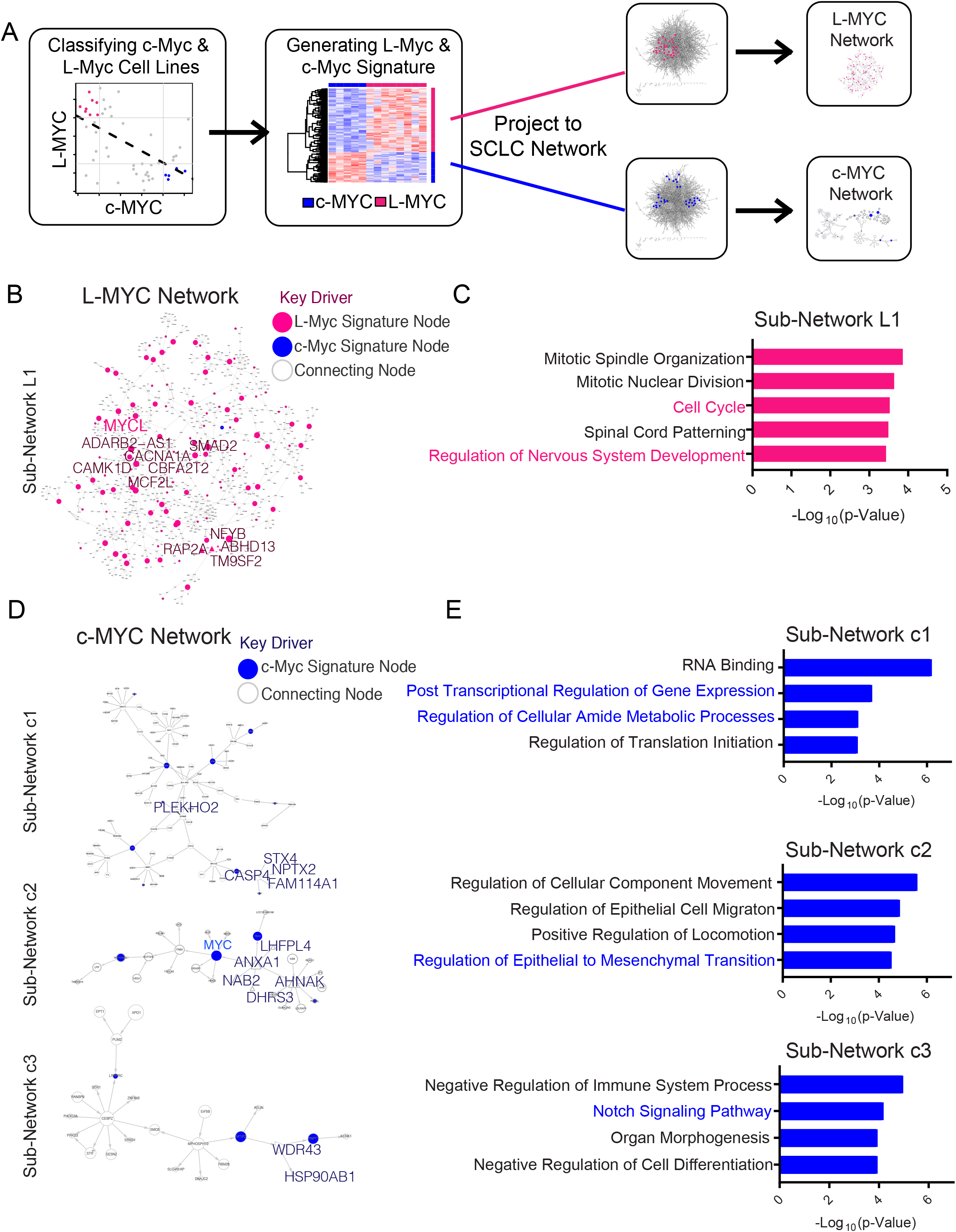
Bayesian network analysis reveals unique L-Myc and c-Myc networks associated with distinct biological processes. A. Schematic of workflow to use SCLC Bayesian causal gene regulatory network to identify networks involving c-Myc and L-Myc. B. L-Myc sub-network showing directionality and association of genes when L-Myc gene signature (Figure S1B) is projected to SCLC Bayesian network. Circles colored in pink represent nodes from L-Myc gene signature. Size of pink circles is directly proportional to the number of outgoing nodes. Nodes indicated in larger text are Key drivers of the sub-network (Table S1). C. Gene Ontology analysis for L-Myc neighbor subnetwork. Enriched functions for these genes are identified based on hypergeometric test against GO terms. D. Three c-Myc sub-networks showing directionality and association of genes when c-Myc associated gene signature (Figure S1B) is projected to SCLC Bayesian network. Circles colored in blue represent nodes from c-Myc gene signature. Size of blue circles is directly proportional to the number of outgoing nodes. Nodes indicated in larger text are Key drivers of the sub-network (Table S2). E. Gene Ontology analysis for corresponding c-Myc neighbor subnetwork. Enriched functions for these genes are identified based on hypergeometric test against GO terms.

To explore the sub-networks associated with L-Myc, we projected the genes upregulated in the *MYCL* expressing subset onto the network and collected all nodes within two layers from them (see Methods). We identified one large closed sub-network (L1, Figure 1B) comprising of 959 gene nodes that included 120 of 310 genes from the L-Myc signature. To identify master regulators of the L-Myc subnetwork, we performed Key Driver Analysis (see Methods) that revealed 13 statistically significant genes (Table S1). Gene ontology analysis of this L-Myc subnetwork revealed enrichments of two biological processes: cell cycle progression and neuronal development (Figure 1C). These two pathways have been previously implicated as core descriptors of classical SCLC (*2*). On the other hand, when we projected the c-Myc signature onto the network, the c-Myc network was organized into three unique sub-networks c1, c2 and c3, comprised of 95, 29 and 25 gene nodes, respectively (Figure 1D). Key Driver Analysis of all three sub-networks revealed 15 statistically significant regulators (Table S2). Sub-network c1 was enriched for canonical c-Myc functions in transcriptional, translational and metabolic regulation (Figure 1E) (*19*). c-Myc sub-network c2 was enriched for Notch signaling pathway (Figure 1E), which has been implicated to mediate a transition from neuroendocrine to non-neuroendocrine fate of tumor cells in a murine SCLC model (*26*). Finally, sub-network c3 was enriched for pathways in epithelial-to-mesenchymal transition (Figure 1E); that has been shown to be relevant in the neuroendocrine-low subtype (*27*). These functional pathways enriched in subnetworks c2 and c3 imply previously uncharacterized role of c-Myc in lineage state determination in SCLC.

Together, these findings suggest that c-Myc and L-Myc are associated with different molecular mechanisms that have been implicated in SCLC biology. Specifically, it implicates c-Myc is associated with neuroendocrine-low differentiation state in addition to its canonical oncogenic functions in SCLC, and by contrast, L-Myc is associated with classic neuroendocrine state of SCLC.

### c-Myc and L-Myc driven SCLCs exhibit distinct chromatin states to exert differential transcriptional programs

Next, we sought to determine whether the distinct c-Myc and L-Myc networks are associated with their distinct cistromes. To examine the role of each Myc family member, we selected representative cell lines for c-Myc (NCI-H82, NCI-H524, NCI-H2171 and NCI-H2081) and L-Myc (CORL-88, NCI-H1963 and NCI-H209) based on the classification described in (Figure S1A) and confirmed protein expression for these transcription factors (Figure 2A). In order to define open regulatory elements potentially regulated by c-Myc and L-Myc, we performed the assay for transposase-accessible chromatin-sequencing (ATAC-seq) on the representative cell lines and filtered identified accessible chromatin regions with motif matrices for Myc (E-box) referring to as Myc accessible regions hereafter (Figure 2B). Of note, inferred Myc accessible regions reasonably recapitulated c-Myc binding sites previously determined by c-Myc chromatin-immunoprecipitation in NCI-H2171 cells (*28*) (Figures S3A and S3B). This approach eliminates confounding factors from antibody-dependent sensitivity and specificity that are intrinsic to ChIP based assays to compare cistromes of two distinct proteins.

**Figure 2:**
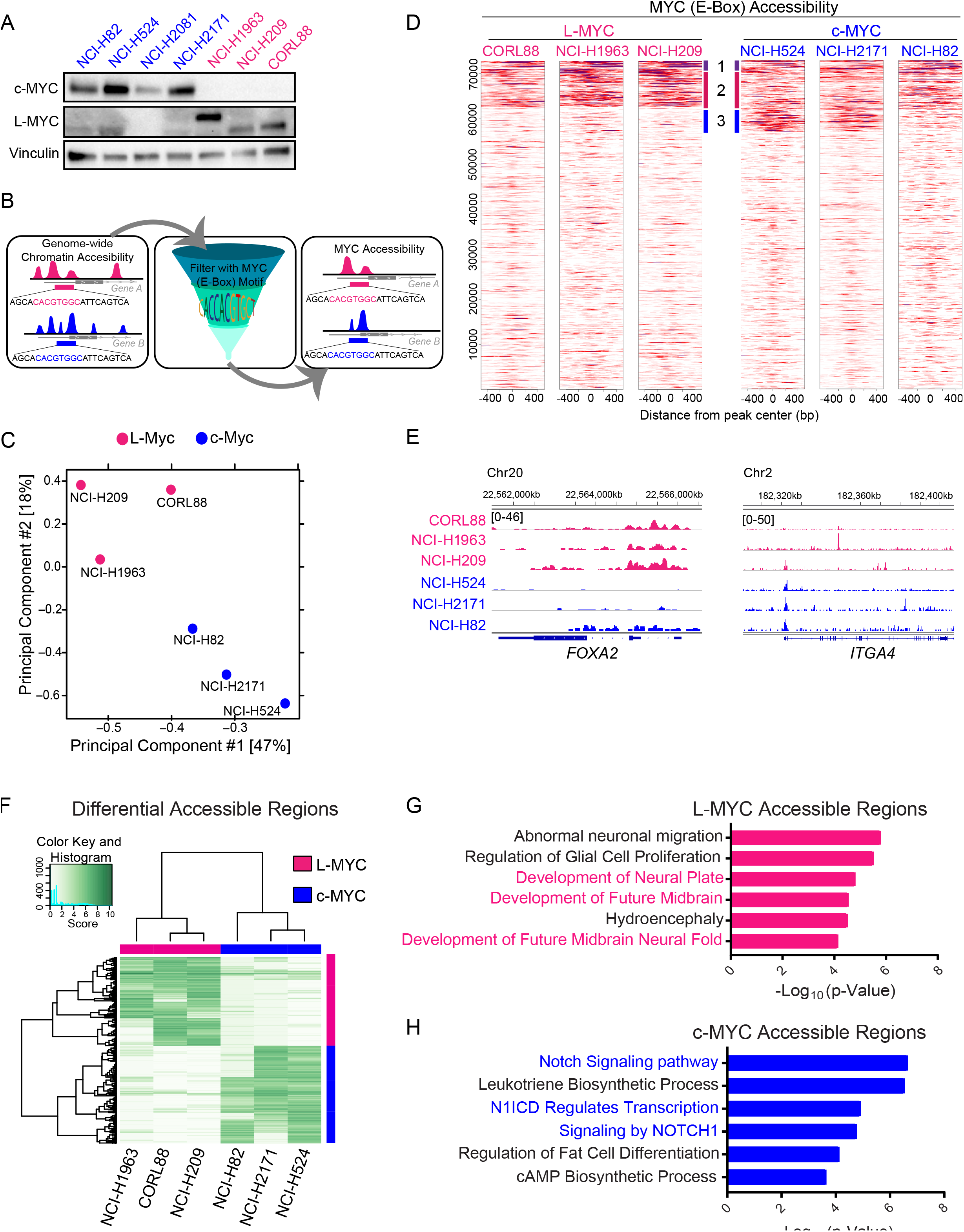
c-Myc and L-Myc driven SCLCs exhibit distinct chromatin states to regulate distinct transcriptional programs. A. Protein expression of c-Myc and L-Myc as well as vinculin as a loading control in a panel of representative SCLC cell lines (blue: c-MYC-classified lines NCI-H82, NCI-H524, NCI-H2171, pink: L-MYC-classified lines NCI-H209, NCI-H1963, CORL88) B. Schematic showing the analytic approach to define Myc accessible regions. C. Principal component analysis of open chromatin regions with E-Box motif in six SCLC cell lines. Each dot represents a SCLC cell line that is colored based on Myc status (blue: c-MYC-classified cell lines, pink: L-MYC-classified cell lines). D. Each heatmap depicting global Myc (E-box) accessibility in individual SCLC cell lines at 72,833 combined peaks. ATAC-seq signal intensity is shown by color shading. E. Genome browser view tracks of ATAC-seq signals in at the *FOXA2* and *ITGA4* loci (blue: c-MYC-classified cell lines, pink: L-MYC-classified cell lines). F. Heatmap showing 2,808 differentially accessible regions (fold change ≥ 5 and FDR ≤ 0.05) between 3 L-Myc cell lines shown in pink and 3 c-Myc cell lines shown in blue. G. Enriched ontology by GREAT analyses for regions differentially accessible in L-Myc classified cells. H. Enriched ontology by GREAT analyses for regions differentially accessible in c-Myc classified cells.

Principal component analysis of peaks containing the E-box motif supported cell line classification according to Myc status (Figure 2C). Additionally, a correlation matrix of pairwise Pearson correlation coefficient of Myc accessibility signal captured distinct clusters for the c-Myc and L-Myc classified cell lines (Figure S3C). We observed a fraction of peaks that overlap between c-Myc classified and L-Myc classified cells (cluster 1; Figure 2D), suggestive of common functional binding that the Myc family members share. We also found distinct groups of peaks with higher signal intensity unique to L-Myc classified cells (cluster 2; Figure 2D) and in c-Myc classified cell lines (cluster 3; Figure 2D), suggesting distinctive DNA binding profiles for L-Myc and c-Myc. We observed preferential accessibility at distinct genetic loci in L-Myc cell lines when compared to c-Myc cell lines (Figure 2E). This included augmented accessibility at lung and neuron development factor *FOXA2* (*29,30*) in L-Myc cell lines and cell surface adhesion factor *ITGA4* in c-Myc cell lines (Figure 2E). Of note, the chromatin landscape of NCI-H82 appeared to be intermediate to the L-Myc and c-Myc profiles. Together, these findings indicate that c-Myc and L-Myc impart differential transcriptional programs.

To gain insights into the unique biological processes c-Myc and L-Myc may impart, we analyzed differentially enriched Myc accessible sites comparing between c-Myc cell lines and L-Myc cell lines. We identified 2,808 differentially accessible regions; 1,235 peaks enriched in L-Myc classified cell lines and 1,573 peaks enriched in c-Myc classified cell lines (Figure 2F). Next, we performed GREAT (Genomic Regions Enrichment of Annotations) analysis (*31*) on the differentially accessible peaks. Unique L-Myc accessible sites are enriched for neuronal pathways such as glial cell proliferation, development of the neural plate, neural fold and future midbrain (Figure 2G). By contrast, unique c-Myc accessible sites are enriched for pathways involved in Notch signaling (Figure 3H). While specific Notch signaling activity in *MYC*-amplified SCLCs has not been previously noted, there have been previous reports showing Rest as a target of the Notch signaling pathway to suppress the expression of neuronal genes (*20,26*).

These findings are consistent with the observations from the Bayesian network analysis providing additional evidence to suggest distinct transcriptional programs in c-Myc and L-Myc driven SCLCs.

### c-Myc and L-Myc expression is associated with distinct lineage state markers

A recent review synthesized SCLC profiling studies to identify four molecular subtypes, where two of these subtypes are classified as neuroendocrine SCLC that are driven by either ASCL1 or NeuroD1, and the other two are classified as non-neuroendocrine driven by YAP1 or POU2F3 (*6*). Given our findings from the Bayesian network and Myc accessibility profiling, that c-Myc and L-Myc regulate transcriptional programs associated with distinct lineage determining programs, we sought to investigate the relationship between Myc family members and the proposed SCLC master regulator classifiers.

Myc accessibility profile at the individual locus level of our c-Myc and L-Myc classified cell lines for these four factors revealed augmented accessibility in the c-Myc classified cell lines at the *NeuroD2* locus in contrast to L-Myc classified cell lines that had augmented accessibility at the *ASCL1* locus (Figure S3A, S3B). We observed no differential or preferential accessibility signal in both c-Myc classified and L-Myc classified cell lines at *YAP2* and *POU2F3* loci likely reflecting that these cell lines do not represent these subtypes (Figure S3C, S3D). Protein expression of these lineage state markers showed that our L-Myc classified cell lines had exclusive expression of ASCL1 while the c-Myc classified cell lines exclusively expressed NeuroD1, consistent with the chromatin profile. (Figure S3E).

To understand how c-Myc and L-Myc expression correlated with the four molecular subtypes in CCLE cell lines, we examined the expression of c-Myc and L-Myc in each of the molecular subtypes as classified in Rudin et al., 2019. We observed that the non-neuroendocrine: POU2F3 and YAP1 as well as NeuroD1 subtypes had higher expression of c-Myc (Figure S3F). On the other hand, L-Myc expression was enriched in the neuroendocrine high ASCL1 subtype (Figure S3G).

We further sought to reproduce the classification with data from primary tumors (*20*) and CCLE cell lines with additional data from 77 primary tumors (*23*) (Figure 3A, Table S4). Of note, while our unsupervised clustering was in general agreement with the published classification (*6*), the ASCL1 subtype cluster revealed heterogeneity particularly of primary tumors with the dual expression of ASCL1 and other lineage factors that may indicate lineage transition or intra-tumor heterogeneity (Figure 3A). In this classification, we found that L-Myc expression was higher in tumors classified as ASCL1 or NeuroD1 subtype when compared to the non-neuroendocrine subtype, thus indicating unique expression of L-Myc in neuroendocrine SCLC (Figure 3B). On the other hand, the expression of c-Myc was higher in tumors and cell lines clustered in the NeuroD1 subtype and non-neuroendocrine low: POU2F3 and YAP1 as compared to those classified as ASCL1 (Figure 3C), suggesting a negative relation between ASCL1 and c-Myc. Indeed, we found the expression of c-Myc is anti-correlated with ASCL1 (r = −0.53) across the combined primary tumor and cell line datasets (Figure 3D). Together, these data imply the dichotomy of c-Myc and L-Myc expression in SCLC.

**Figure 3:**
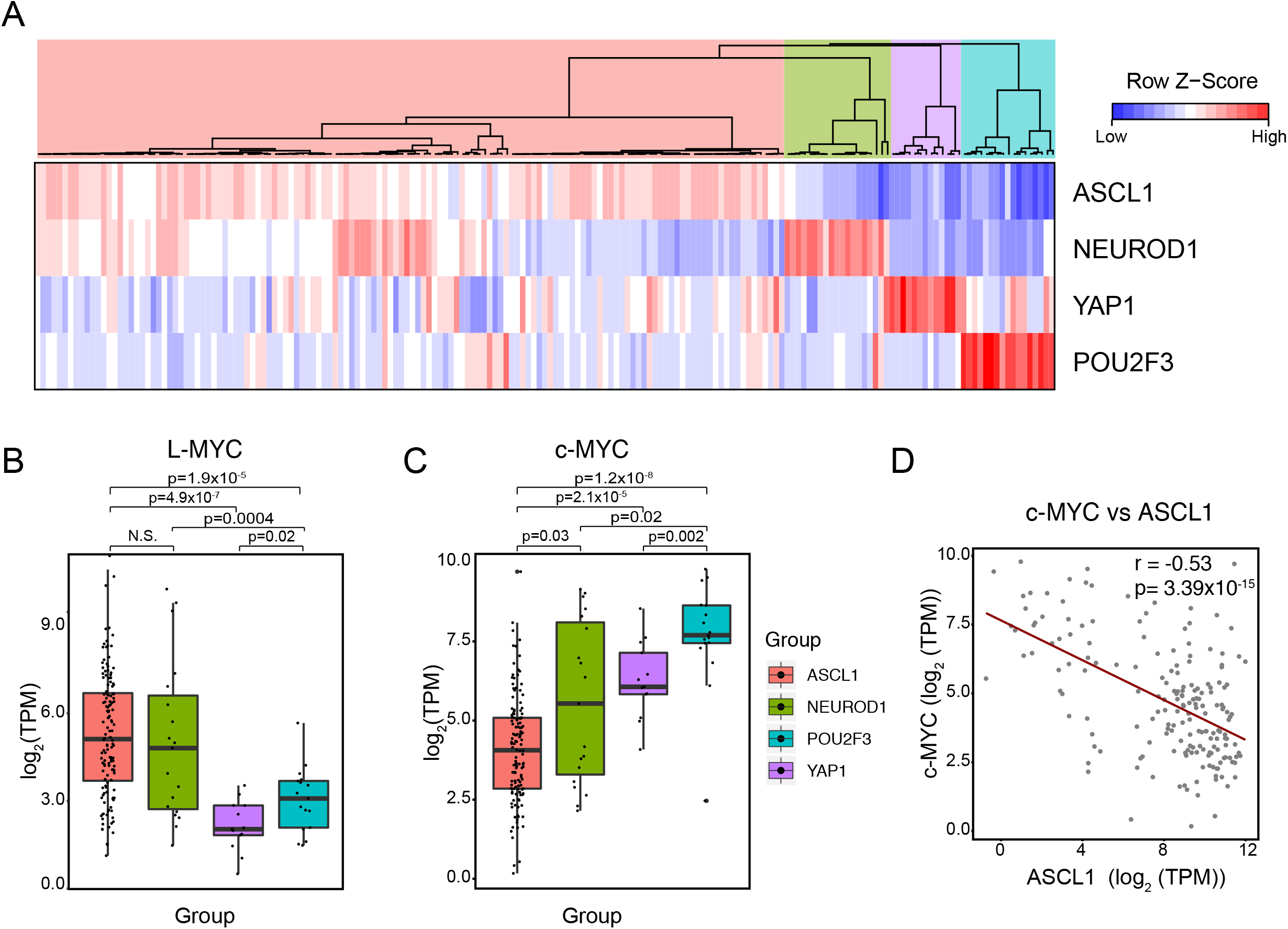
c-Myc and L-Myc expression is associated with distinct molecular subtype markers. A. Unsupervised clustering of 137 primary tumors and 51 CCLE cell lines using the expression of ASCL1, NeuroD1, POU2F3 and YAP1 distinguishing four molecular subtypes. (Table S3) B. Box plot showing mRNA expression of L-Myc across four molecular subtypes for combined primary tumor and cell line classification described in Figure 3A. C. Box plot showing mRNA expression of c-Myc across four molecular subtypes for combined primary tumor and cell line classification described in Figure 3A. D. Scatter-plot with best-fit line showing negative correlation between c-Myc and ASCL1 expression for combined primary tumor and cell lines. Pearson’s correlation coefficient (r) = −0.53, *P* <0.01 t-test.

### L-Myc induces a neuronal state but fails to transition to ASCL1 expressing SCLC

Given that L-Myc and c-Myc expression are enriched in distinct subtype(s) and they regulate programs associated with its respective subtype, we hypothesized that part of the role Myc family members play is to serve as a lineage factor. To test the potential of L-Myc to establish the neuroendocrine lineage, we modified a c-Myc expressing NeuroD1 classified cell line NCI-H82 by exogenous overexpression of L-Myc and CRISPR-Cas9 mediated deletion of c-Myc (Figure 4A). We found c-Myc expressing cell line (NCI-H82) was able to tolerate the overexpression of L-Myc (Figure 4B). Cells engineered to express both L-Myc and c-Myc exhibited a modest increase in proliferation rates compared to parental cells expressing only c-Myc (Figure 4C). We further demonstrated that replacement of c-Myc with the expression of L-Myc allowed the cells to retain proliferative potential in the absence of c-Myc in NCI-H82 (Figure 4C). This indicates redundant roles among the two Myc family members in regards to maintaining cell viability.

**Figure 4:**
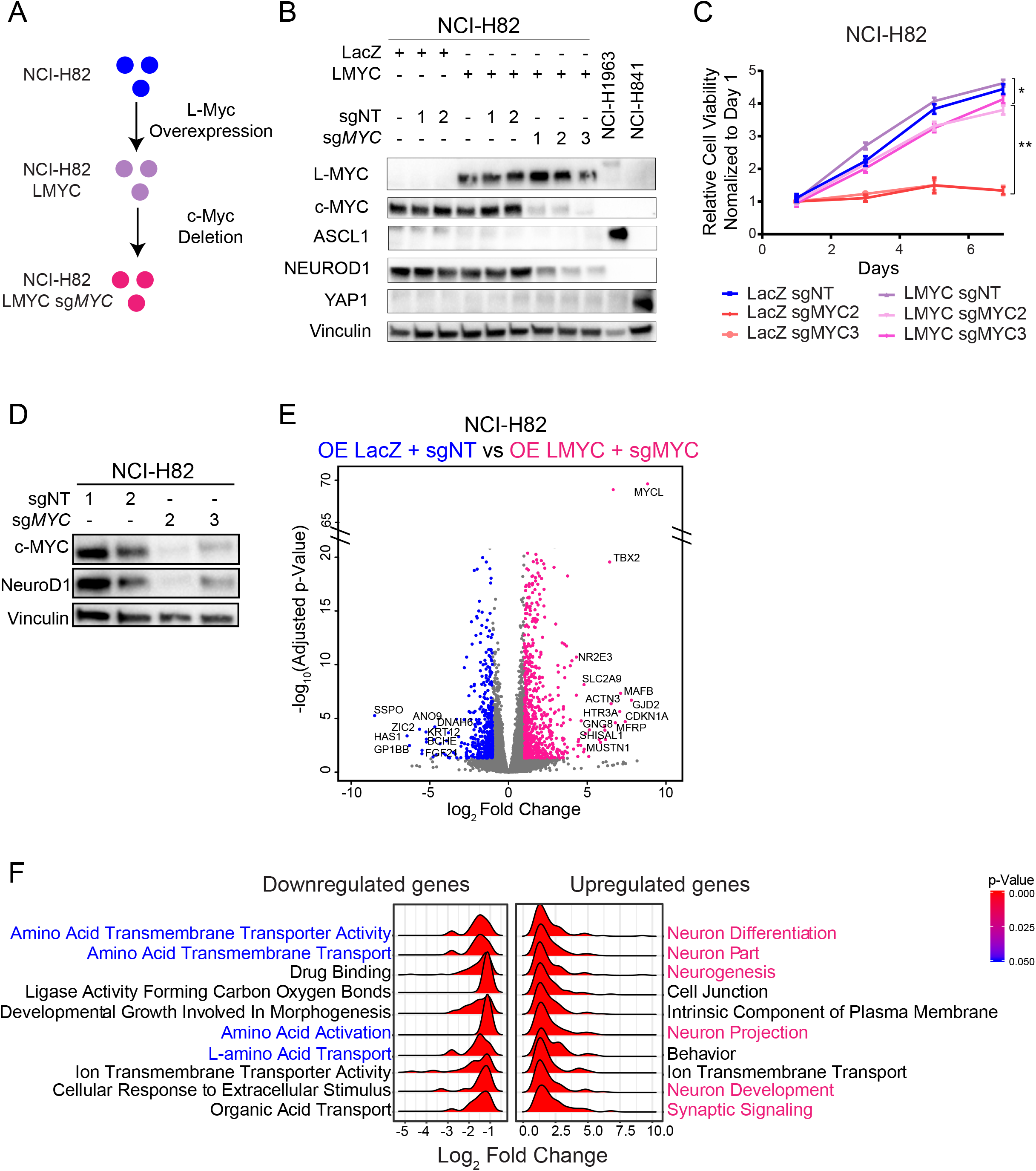
c-Myc is required to maintain NeuroD1-positive lineage state. A. Schematic showing the workflow to genetically engineer replacement of c-Myc with L-Myc in NCI-H82. B. Protein expression of L-Myc, c-Myc, ASCL1, NeuroD1 and YAP1 as well as vinculin as a loading control in genetically engineered NCI-H82 cells to replace c-Myc with L-Myc. C. Cell growth curve for NCI-H82 cells with over-expression of L-Myc and genetic deletion of c-Myc. **, *P* <0.01 vs. Lac-Z overexpressed + sgNT NCI-H82 cells, t-test. *, *P* <0.05 vs. Lac-Z overexpressed + sgNT NCI-H82 cells, t-test. D. Protein expression of c-Myc and NeuroD1 with vinculin as loading control in genetically engineered NCI-H82 cells with c-Myc ablation. E. Volcano plot showing 1,213 differentially expressed genes (713 upregulated in pink and 500 downregulated in blue) between control NCI-H82 cells (LacZ-overexpressed cells and LacZ-overexpressed cells with nontarget sgRNA) and engineered NCI-H82 cells (LMYC-overexpressed cells with three sgRNA targeting MYC). F. Ridgeplot showing distribution of fold-change for genes that belong to each enriched gene ontology term for differentially upregulated (right panel) and downregulated genes (left panel).

To inquire their contribution to lineage determination, we evaluated the expression of lineage state markers. Addition of L-Myc neither induced expression of neuroendocrine-marker ASCL1 nor altered the expression of variant state marker NeuroD1 (Figure 4B). By contrast, the replacement of c-Myc with L-Myc led to the downregulation of NeuroD1. (Figure 4A). Without L-Myc replacement, consistent with previous findings (*32*), c-Myc was essential for survival in NCI-H82 (Figure 4C), thus precluding us from evaluating converted lineage state. However, during the short window of survival, c-Myc ablation led to significant downregulation of NeuroD1 suggesting a role of c-Myc on maintaining expression of lineage state marker (Figure 4D). The lack of expression of neuroendocrine-low lineage state marker YAP1 in these ASCL1- and NeuroD1-negative cells also indicates a unique lineage negative state for these cells (Figure 4B).

Previous literature showed that deletion of *Mycl* (L-Myc) in the classical murine SCLC model resulted in tumors with mixed and NSCLC morphology (*15*) in contrast to the c-Myc expressing SCLC with ‘variant’ morphology (*9,17*). Therefore, to investigate L-Myc’s contribution to SCLC histology, we implanted genetically engineered NCI-H82 cells into immunodeficient mice. Histological analysis of NCI-H82 expressing L-Myc and NCI-H82 L-Myc ablated with c-Myc xenografts revealed morphology similar to the control NCI-H82-LacZ+sgNT cells with polygonal cells, large nucleoli and moderate amounts of cytoplasm representative of ‘variant’ SCLC consistent with previous reports (*9*) (Figure S4A). The insufficiency of L-Myc to induce histological trans-differentiation may explain the low frequency of transition from variant to classical SCLC (*26*).

To investigate the state of these lineage-marker negative cells, we performed transcriptomic profiling. Gene ontology analysis of 235 significantly upregulated genes by L-Myc in the presence of c-Myc in NCI-H82 revealed enrichment for neurogenesis and neuronal associated pathways (Figures S4B & S4C), suggesting that L-Myc imparts neuronal pathways while not fully inducing ASCL1-lineage state. Ablation of c-Myc further upregulated 170 genes compared to the previous condition, which are enriched for neuronal structure, suggesting the cells are further committed to a neuronal state (Figure S4D and S4E). A direct comparison of the transcriptome control NCI-H82 cells with lineage negative-NCI-H82 cells, revealed 713 significantly upregulated genes enriched for neuronal associated pathways and 500 downregulated genes enriched for canonical Myc function (Figure 4E and 4F). Collectively, the data suggest that replacement of c-Myc with L-Myc induces a profile that represents more neural state but not fully transdifferentiates to ASCL1-positive state and remains to be negative for lineage markers used in the current consensus molecular classification (*6*).

### Aurora Kinase A Inhibition sensitivity is altered with change in Myc status

Previous findings suggest that *MYC* (c-Myc)-driven SCLC is more responsive to Aurora kinase A inhibition (*17,33*). We confirmed previous findings that *MYC*-amplified cell lines exhibit increased sensitivity and *MYCL*-amplified cell lines are resistant to alisertib (Figure S5A). Alisertib treatment of the NCI-H82 cells expressing both c-Myc and L-Myc revealed no changes in sensitivity to the drug nor did changes in expression of the variant state marker. However, the cells that replaced c-Myc with L-Myc became resistance to aurora kinase inhibition (Figure S5B). On the other hand, overexpression of c-Myc increased sensitivity at lower concentrations of the drug (Figure S5C). This suggests either alisertib sensitivity is attributable to c-Myc driven lineage state of the cells, or difference in molecular interaction of Aurora kinase A with each Myc protein.

### c-Myc causes loss of classical neuroendocrine SCLC features

The dependency of lineage state marker NeuroD1 on c-Myc led us to hypothesize that c-Myc exerts a role in addition to its oncogenic role to regulate and establish a variant differentiation state. To this end, we genetically engineered a neuroendocrine-high *MYCL*-amplified NCI-H1963 with inducible exogenous overexpression of c-Myc (Figure 5A). Of note, exogenous expression of c-Myc led to downregulation of L-Myc in NCI-H1963.

**Figure 5:**
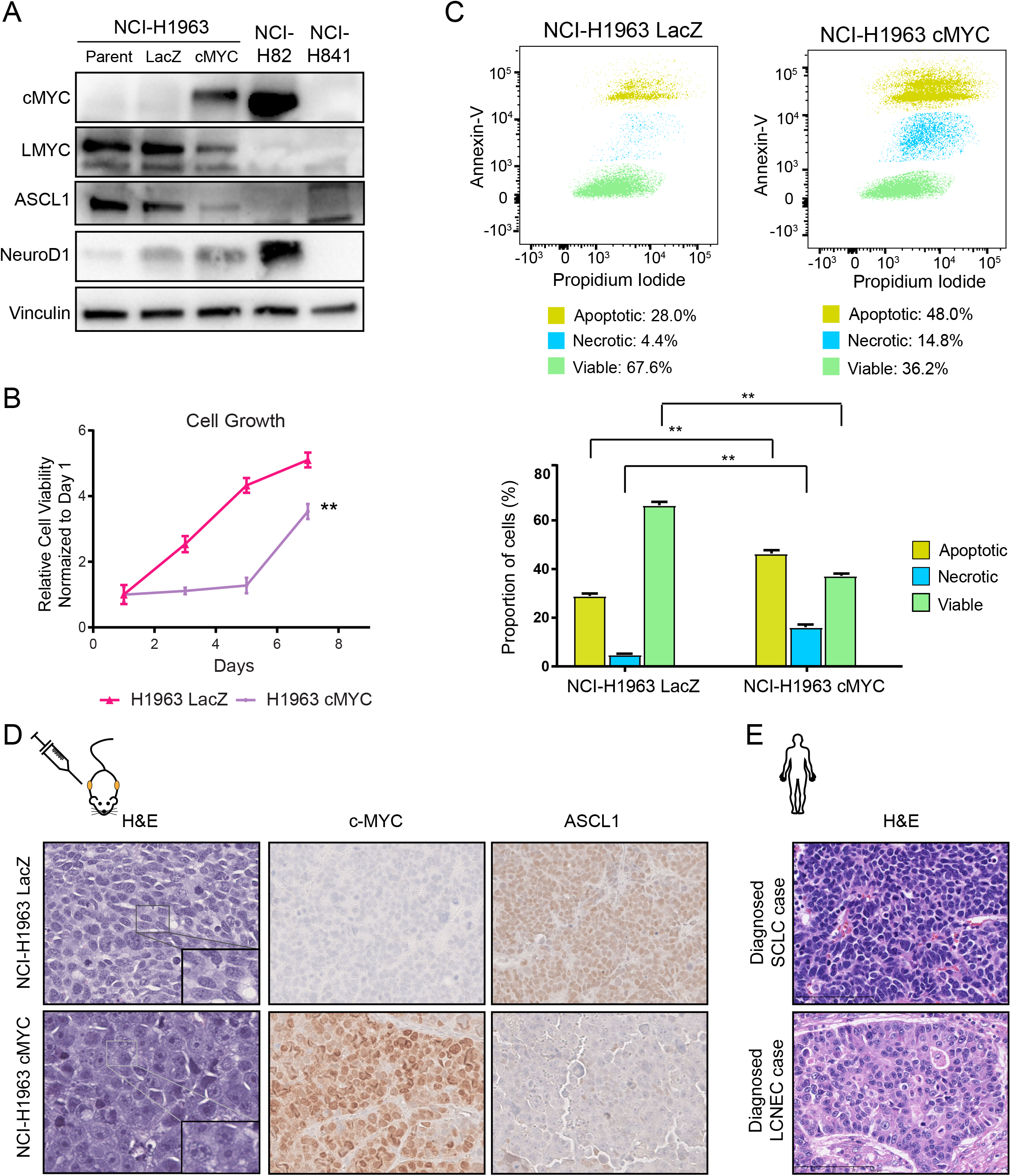
c-Myc expression in ASCL1-subtype SCLC induces trans-differentiation to histologically distinct NeuroD1-positive state. A. Protein expression of c-Myc, L-Myc, ASCL1, and NeuroD1, as well as vinculin as loading control in genetically engineered NCI-H1963 cells. B. Cell growth curve for NCI-H1963 cells with over-expression of c-Myc or control Lac-Z. Mean ± SD of five technical replicates. **, *P* <0.01 vs. Lac-Z overexpressed NCI-H1963 cells, t-test. C. Scatter plot showing Annexin-V staining for NCI-H1963 cells with over-expression of c-Myc (right panel) or over-expression of Lac-Z as control (left panel) with proportions of viable, necrotic and apoptotic cells. Bar chart shows mean ± SD of three biological replicates. ** *P* <0.01, t-test. D. Hematoxylin-eosin staining and immunohistochemical staining of c-Myc and ASCL1 in LacZ-overexpressed (top panel) or c-Myc-overexpressed (bottom panel) NCI-H1963 xenograft. Original images, x40. Inset shows cell morphology. E. Hematoxylin-eosin staining of typical SCLC and LCNEC primary human tumors. Original images, x400.

When c-Myc expression is introduced in NCI-H1963, we observed an initial phase of growth suppression (Figure 5B). This observation stands in contrast to the established role of c-Myc to promote cell cycle and cell proliferation (*34*). The growth suppression is unlikely to be due to oncogene induced senescence as these cells stained negative for beta galactosidase (Figure S6A) and are deficient for Rb1 and p53. We investigated the cell cycle dynamics in their growth suppressive phase, and found that initial c-Myc expression (day 2) modestly increased the proportion of cells in S and G2/M phase but did not significantly alter the distribution of cells or induce cell cycle arrest to explain slower cell growth (Figure S6B). Alternatively, we investigated the proportion of Annexin-V positive cells on day 2 and found c-Myc overexpression increased the proportion of necrotic and apoptotic cells as compared to control cells (Figure 5C).

We hypothesized that the cell death was a consequence of incompatibility of the neuroendocrine differentiation state with c-Myc expression and that the persisting cells eventually grown out were cells that tolerated the expression of c-Myc, potentially by switching into a differentiation state compatible with c-Myc expression (Figure 5B). Therefore, we investigated the expression of lineage state markers and found that the overexpression of c-Myc downregulated ASCL1 and upregulated NeuroD1 in NCI-H1963 cells (Figure 5A), indicating the trans-differentiation to NeuroD1-subtype of SCLC and that c-Myc exerts a role in dictating a distinct NeuroD1-positive neuroendocrine differentiation state.

To investigate if the trans-differentiation between molecular subtypes was accompanied by histological switch *in vivo,* we injected NCI-H1963 cells genetically engineered with stable c-MYC expression. We observed that NCI-H1963 overexpressed with LacZ (control cells) had oval and elongated nuclei with a high nuclear:cytoplasm ratio, smooth granular chromatin, no nucleoli consistent with classical SCLC histology (Figure 5D, Figure S6C), similar to the histology that would be typically diagnosed with SCLC in primary human tumors (Figure 5E). Histological analysis of xenografts of NCI-H1963 overexpressed with c-Myc showed presence of nucleoli and polygonal shaped larger cells, reminiscent of variant SCLC histology as defined by (*9*)(Figure 5D, Figure S6C). The histopathology observed here, according to the current WHO classification, would likely be recognized as either purely LCNEC or “combined SCLC” that includes a component of large-cell neuroendocrine carcinomas (LCNEC), potentially reflective of the discrepancies between clinical practice and experimental findings (Figure 5E) (*10,35*). Together, these results suggest that c-Myc expression drives the transition from ASCL1-positive state characterized by classical SCLC histology to NeuroD1-positive state SCLC characterized by ‘variant’ SCLC/LCNEC histology.

### c-Myc mediates trans-differentiation through REST

We sought to investigate the mechanism through which c-Myc mediates transdifferentiation to the ASCL1-negative/NeuroD1-positive state. To this end, we profiled the accessible chromatin profile of trans-differentiated NCI-H1963. Filtering the regions to reflect regions with Myc (E-box) motif matrices reflected a common pattern (clusters 1 to 4). c-Myc overexpression resulted in the gain of additional regulatory regions (cluster 5) and a decrease in accessibility at a fraction of sites in cluster 6 (Figure 6A), revealing an alteration of the Myc (Ebox) cistrome and further suggesting non-redundancy of this family of transcription factors. Next, we sought to investigate how the re-wiring of these regulatory regions compared to c-Myc and L-Myc expressing SCLC. Principal component analysis of Myc accessibility in these genetically engineered cells and c-Myc and L-Myc classified cell lines revealed NCI-H1963 expressing c-Myc clustered closer to c-Myc classified cell lines along Principal Component #1, mapping NCI-H1963 transitioning from L-Myc cistrome towards a c-Myc cistrome (Figure 6B). These data re-iterate c-Myc mediated alteration in regulatory regions that dictate cellular identity.

**Figure 6:**
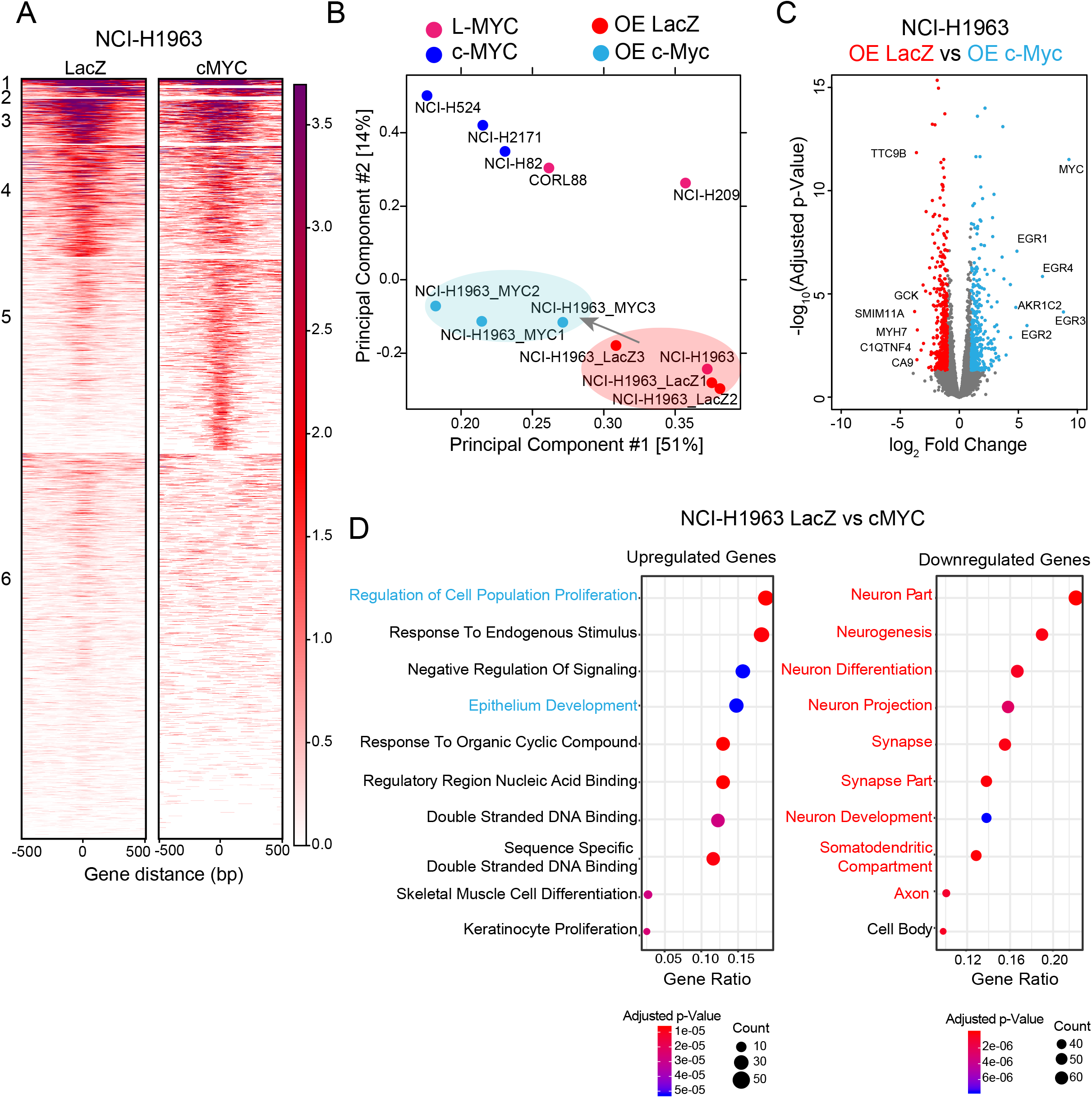
c-Myc rewires Myc-accessible chromatin regulatory elements to regulate distinct lineage associated pathways. A. Each heatmap depicting *k-means* (k =6) clustering of global Myc (E-box) accessibility in NCI-H1963 overexpressing LacZ (left) and NCI-H1963 overexpressing c-MYC (right) at 44,722 combined peaks. ATAC-seq signal intensity is shown by color shading. B. Principal component analysis of open chromatin regions with E-Box motif in six SCLC cell lines in addition to NCI-H1963-LacZ (control) and NCI-H1963-c-Myc in triplicate. Each dot represents a SCLC cell line that is colored based on Myc status (pink: L-MYC, blue: c-MYC, red: NCI-H1963 LacZ, light blue: NCI-H1963 cMYC). C. Volcano plot showing 775 differentially expressed genes (375 upregulated in light blue and 400 downregulated in red) between control NCI-H1963 cells with LacZ-overexpression cells and NCI-H1963 cells with c-Myc-overexpression. D. Dot plot showing gene ontology analysis for differentially upregulated (right panel) and downregulated (left panel) genes.

To identify the transcriptional programs associated with the changes in regulatory regions of the MYC cistrome, we performed RNA-seq on trans-differentiated NCI-H1963. We identified 775 differentially expressed genes (375 upregulated genes downregulated and 400 downregulated genes) (Figure 6C). Gene ontology analysis for upregulated genes revealed an enrichment for epithelium development in addition to canonical c-Myc functions including regulation of cell proliferation and regulatory region nucleic acid binding (Figure 6D, right panel) while gene ontology analysis for downregulated of genes revealed an enrichment for neuronal associated genes (Figure 6D, left panel). These findings suggest c-Myc expression in ASCL1 expressing SCLC induces a transition to a more epithelial state and are consistent with the loss in neuroendocrine lineage state marker (Figure 5A), but did not reveal further specific mechanistic insights.

Previous works have shown the negative regulation of ASCL1 by Notch signaling during lung development (*36*) and activation of Notch signaling in SCLC suppresses ASCL1 expression (*20, 37, 38*). Additionally, our SCLC transcriptional network analysis and Myc accessibility data revealed Notch pathway activation preferentially in c-Myc expressing tumors and cell lines (Figures 1E and 3C). Although we did not observe an enrichment for an active Notch signaling signature amongst the upregulated genes, when we investigated the expression of known Notch signaling pathway targets in trans-differentiated NCI-H1963, we identified Notch-target RE-1 Silencing transcription factor (REST) that is a transcriptional repressor of neuronal genes and ASCL1 (*26*) as one of the genes that were induced upon c-Myc expression (Figure 7A) and we confirmed this with protein expression (Figure 7B). Therefore, we sought to investigate the c-Myc–Notch–Rest axis in mediating trans-differentiation, and pharmacologically inhibited Notch1-4 using a γ-secretase inhibitor (GSI), DBZ (*39*) in combination with c-Myc expression in NCI-H1963 cells. Inhibition of Notch signaling, did not alter expression of Rest and ASCL1 (Figure 7C). Next, to investigate if Rest is required for c-Myc mediated suppression of ASCL1, we used pharmacological inhibition of Rest (*40*) in c-Myc over-expressed NCI-H1963 cells. We confirmed treatment with Rest inhibitor (X5050) led to a decrease in Rest protein expression. Surprisingly, simultaneous suppression of Rest and expression of c-Myc did not result in decrease in the level of ASCL1 (Figure 7D). These data suggest a novel Notch-independent, c-Myc mediated activation of Rest and NeuroD1 and Rest mediated-suppression of ASCL1.

**Figure 7:**
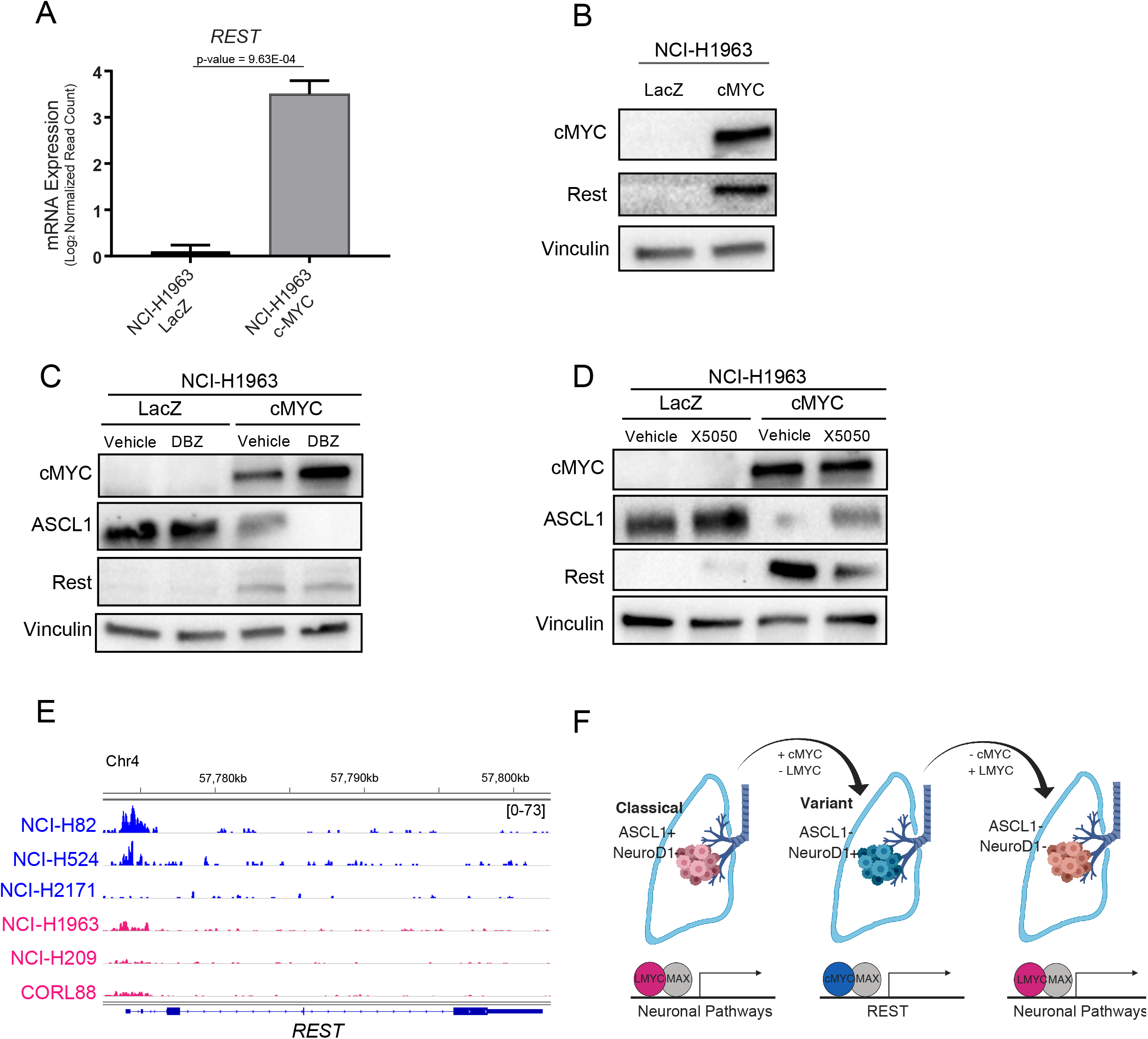
c-Myc induces neuronal repressor REST to mediates lineage conversion. A. Expression of *REST* in NCI-H1963 cells with LacZ over-expression and NCI-H1963 with c-Myc overexpression (n =3). B. Protein expression of c-Myc, and Rest, as well as vinculin as loading control in genetically engineered NCI-H1963 cells. C. Protein expression of c-Myc, L-Myc, ASCL1, and Rest, as well as vinculin as loading control in genetically engineered NCI-H1963 cells treated with vehicle (DMSO) or 0.5 μM of a Notch inhibitor (DBZ) for 48 hours. D. Protein expression of c-Myc, L-Myc, ASCL1, and Rest, as well as vinculin as loading control in genetically engineered NCI-H1963 cells treated with vehicle (DMSO) or 10 μM of a Rest inhibitor (X5050) for 48 hours. E. Schematic showing the role of c-Myc and L-Myc in SCLC lineage specification

## Discussion

In this study, we showed a novel functional distinction between c-Myc and L-Myc in SCLC, in contrast to the prevailing notion that molecular function of Myc family members are similar to each other (*15*). We found c-Myc is associated not only with canonical roles including transcriptional and translational regulation, but also with lineage associated pathways such as Notch signaling and epithelial-mesenchymal transitions. Our findings are supported by a previous report that described active Notch signaling negatively regulating neuroendocrine lineage by suppressing the expression of *ASCL1* (*26*) as well as a study that showed mesenchymal-like features in neuroendocrine-low sub-type of SCLC (*27*). On the other hand, we found L-Myc is associated with neuronal pathways implicating their relevancy in determining the classical neuroendocrine state of SCLC. These findings are consistent with previous observation showing a correlation of expression between L-Myc and neuronal proteins (*41*). Taken together, contrary to the notion that Myc family members, c-Myc in particular, exclusively act as oncogenes and general amplifiers of expression and metabolism, our data suggest that they additionally exert lineage defining transcriptional programs (*28*). We propose that the Myc family members regulate a defined set of transcriptional programs that are essential in SCLC lineage subtype determination. Of note, although these biological pathways associated with these factors have been shown to be relevant for lineage maintenance and determination in distinct subtypes of SCLC, their association with Myc family members has not been described before.

We found L-Myc failed to induce the expression of a neuroendocrine lineage state marker ASCL1, suggesting its inability to fully control trans-differentiation from neuroendocrine-low/variant state to neuroendocrine high/classical state. However, based on previous work suggesting requirement of L-Myc to drive tumors of neuroendocrine origin in SCLC GEMM (*15*) and our findings that revealed L-Myc regulating and inducing neuronal development pathways, L-Myc likely plays roles in lineage state maintenance and/or is more compatible with the neuroendocrine lineage.

We also report a previously undescribed function for c-Myc in lineage maintenance in NeuroD1-positive SCLC. c-Myc expression was incompatible with ASCL1-positive SCLC and induces trans-differentiation to NeuroD1-positive state revealing the role for c-Myc in regulating lineage plasticity. We show that this trans-differentiation between molecular subtypes of SCLC is accompanied by alterations in the transition of histopathology in SCLC from classical SCLC features to variant histopathology. These data reflect that the variant histopathology, that would typically be classified as “combined” SCLC or LCNEC per the current guidelines of WHO, can indeed be derived from SCLC. This itself has significant clinical implication. This phenotype is reflective of biology distinct from classical SCLC and may warrant distinct therapeutic strategies.

We identified that the trans-differentiation is mediated independent of Notch signaling but through the direct activation of a Notch signaling target, Rest, a transcriptional repressor. These findings may be clinically important when evaluating the rationale for targeting SCLC by activating Notch signaling with drugs including LSD1 and KDM5A (*37,38*). Our data suggests that downstream actors of Notch signaling may induce the trans-differentiation to NeuroD1positive or variant subtype. This may not be a favorable outcome since the variant subtype has been reported to be more frequent in tumors after initial response to chemotherapy (*9,17*). We found that the altered differentiation state with the gain of L-Myc and loss of c-Myc cells conferred resistance to Aurora Kinase A inhibition and in contrast the trans-differentiation as a result of c-Myc gain and loss of L-Myc cells conferred sensitivity to Aurora Kinase A inhibition. The differential sensitivity associated with distinct lineage state with Myc family member status as a biomarker could translate to significant clinical implications in stratification of patients for targeted therapy.

SCLC is a recalcitrant disease typically characterized by neuroendocrine differentiation, nonetheless approximately 10-20% of SCLCs may lack expression of diagnostic neuroendocrine markers (*35*). Here we report, the plasticity between these histological subtypes and molecular subtypes is regulated by c-Myc and L-Myc. The role in lineage determination and maintenance for this family of transcription factors is striking since the Myc family has historically been grouped as general oncogenes. Our data suggest that the role of Myc family in SCLC tumorigenesis could be re-defined. This will enable us to categorize these subtypes to develop effective therapies to combat this highly lethal disease.

## Methods

### CCLE RNA-seq analysis

RNA-seq gene expression data and SNP-array copy number data for *MYC, MYCL* and *MYCN* were downloaded from (CCLE) web site. Log_2_(RSEM (transcripts per million) +1) values for mRNA expression and log2(inferred copy number/2) values for copy numbers were used to depict a heatmap for a 49 SCLC cell lines. *MYC* classified cell lines were further refined with expression cut-off for *MYC* at log2 RSEM > 8 and expression of *MYCL* at log2 RSEM < 2. For MYCL classified cell lines expression cut-off was set at *MYCL* log2 RSEM > 8 and expression of *MYC* log2 RSEM < 2.

### Bayesian Network Analysis

A comprehensive regulatory network was built using the software suite, Reconstructing Integrative Molecular Bayesian Network (RIMBANet) (Zhu et al., 2004) by combining RNAseq data from two independent SCLC primary cohorts (George et al, 2015; Jiang et al., 2016) (detailed in Yoo et al. manuscript in preparation). To identify sub-networks enriched for L-Myc and c-Myc signatures, the gene signature was overlaid on the network and neighboring nodes within two layers for each signature were selected. Then closed subnetwork with more than 25 nodes were collected for L-MYC and c-MYC subnetworks. Then, Key Driver Analysis (*42*) identified master regulators associated with L-Myc and c-Myc signatures in two steps. First, for each node in the network, we compared neighboring nodes within two layers for each signature to obtain a list of potential master regulators whose overlap is statistical significant (Fisher’s exact test (FET) p-value<0.05/(# of nodes)). After sorting candidate genes by corresponding FET p-values, the gene with the strongest p-value was determined as a key regulator. Any candidate regulators in its two-layered neighbors were excluded from the candidate list. The process was iterated throughout the sorted candidate list. Enriched functions were identified using GO terms in Molecular Signatures Database (MSigDB). The significance of the enrichment was determined based on FET p-value considering multiple testing (FET p<0.05/(# of dataset tested)) in each category.

### Cell Lines

All cells were cultured at 37°C in a humidified incubator at 5% CO_2_. SCLC cell lines NCI-H2171, NCI-H82, NCI-H524, NCI-H209, NCI-H1963, CORL88, HCC33, NCI-H69, and NCI-H526 were maintained in RPMI1640 (Gibco), and supplemented with 10% Fetal Bovine Serum (FBS, Sigma) and 1mM Penicillin Streptomycin (P/S, Gibco). HEK293T cells were maintained in Dulbecco’s Modified Eagle’s Medium (DMEM), and supplemented with 10% FBS (Sigma) and 1mM P/S. Cultured cells were regularly tested for mycoplasma using the mycoAlert Detection Kit (Lonza).

### ATAC-seq

ATAC-seq was performed as previously described (*43*). Five thousand cells were harvested and pre-treated with DNase I (Thermo Fisher Scientific) at 37°C for 20 minutes to eliminate DNA contamination from dead cells. The cells were washed in cold Phosphate-buffered saline (PBS) to eliminate traces of DNase. Nuclei were isolated with nuclear lysis buffer (10 mM Tris, 10mM NaCl, 3mM MgCl2 0.1% IGEPAL-630) and centrifuged at low speeds. The nuclei were resuspended in transposase reaction mixture (10mM Tris PH 8.0, 5 mM MgCl2, 10% Dimethlyformamide (DMF), 0.2 mg/ml Tn5 transposase complex). Transposition was carried out at 37°C for 30 minutes followed by DNA purification with DNA Clean and Concentrator-25 (Zymo Research) according to manufacturer’s recommendation. Following purification, library fragments were PCR amplified with Nextera adapter primers. Sequencing was performed at Tisch Cancer Institute sequencing core on the NextSeq500 (Illumina) for 38 nucleotides each from paired ends according to manufacturer’s instructions.

### ATAC-seq data analysis

Illumina sequencing adapter was removed using Cutadapt from raw sequence files in fastq format. The reads were aligned to the hg19 reference genome using Bowtie2 with -k 1 parameters. The aligned reads were used to eliminate PCR duplicates using samtools and filtered off an ATAC blacklist for mitochondrial DNA homologs generated by (*43*). Fragment ends were shifted +4 nucleotides for positive strand and −5 for negative strand to account for distance from Tn5 binding and helicase activity to identify cut sites. Extended Tn5 cut sites were used for peak calling with MACS2 with parameters --nomodel --extsize 100 --shift 50 --nolambda --keep-dup all. Fraction of reads in peaks (FRiP) score was calculated for each sample and was used as a normalization factor for pileup rescaled to a total number of uniquely alignable sequences by WigMath function of Java-Genomic Toolkit. Peaks were filtered by MYC motif using findMotifsGenome function in homer using four motif matrices *MA0058.2 (MAX), MA0059.1 (MAX::MYC), MA0104.4 (MYCN), and MA0147.3 (MYC*). Peaks were quantized using *k*-means clustering (*k*=8) and heatmaps for the peaks filtered with MYC motif were generated using cistrome ((*10,35*)(*11*)) or using *k*-means clustering (*k*=6) and plotHeatmap function of deepTools (*44*) using wiggle files for genetically engineered NCI-H1963. Normalized ATAC-seq signals for each sample was visualized on integrative genome viewer (IGV) genome browser (*45*).

Differentially bound sites comparing three c-Myc classified cell lines vs three L-Myc classified cell lines were identified using R package DiffBind with cutoffs set at fold change ≥ 5 and FDR ≤ 0.05. Functional analysis of the differentially bound regions was performed using Genomic Regions Enrichment of Annotations Tool (GREAT)(*31*).

### Western Blot and Antibodies

Protein lysates were prepared by washing cells with 1 ml of cold PBS and resuspended in lysis buffer (150 mM NaCl, 50 mM Tris-HCl at pH 8.0, 1% NP-40, 0.5% Na deoxycholate, 0.1% SDS, protease inhibitors) for 30 min at 4°C. Lysates were centrifuged at 5°C for 15 minutes at 13,000 rpm to remove insoluble debris. Protein concentrations were quantified using Pierce™ BCA (Thermo Fisher Scientific). Proteins were separated by electrophoresis on a SDS-PAGE gel (BioRad), transferred to a PVDF membrane (Thermo Fisher Scientific) and blocked with 5% milk in Tris-buffered saline with Tween-20 (TBS-T). The membranes were immunoblotted with anti-c-Myc (Santa Cruz Biotechnology), anti-L-Myc (Proteintech), anti-vinculin (Sigma), anti-ASCL1 (BD Biosciences) and anti-NeuroD1(Proteintech), anti-β-Actin antibody (Sigma), or anti-vinculin (Sigma).

### Lentiviral introduction of genes

c-Myc, L-Myc or LacZ open reading frame (ORF) was cloned into pLEX_307 (a gift from David Root, Addgene #41392) using the Gateway^®^ cloning methods according to manufacturer’s recommendations. (Thermo Fisher Scientific). HEK293T cells were seeded in a 10 cm tissue culture dish and incubated at 37°C and 5% CO_2_. At 80% confluency the cells were co-transfected with 10 μg of plasmid constructs, 7.5 μg of psPAX2 (a gift from Didier Trono, Addgene #12260) and 2.5 μg of pMD2.G (a gift from Didier Trono, Addgene #12259) vectors using TransIT-Lenti (Mirus) following manufacturer’s recommendations. At 48h post transfection, virus-containing supernatants were collected, filtered (0.45 μm) and stored at −80°C. Cells were infected with lentiviral supernatant supplemented with polybrene at a final concentration of 8 μg/mL or lentiBlast (OZ BioSciences) at a ratio of 1:1000. Cells were selected with puromycin (1-2 μg/mL for 4-6 days).

### CRISPR-Cas9 genome editing

Cells with stable human codon-optimized S. pyogenes Cas9 expression were generated by infection with the lentiCas9-Blast plasmid (a gift from Feng Zhang, Addgene # 52962). sgRNAs targeting *MYC* and *MYCL* were selected from the Brunello library (*46*). Non-target sgRNA from the Gecko library v2 (*47*) were used as non-target sgRNAs. sgGRNA target sequences are listed in Supplementary Table S3. sgRNAs were cloned using BbsI site downstream of the human U6 promoter in a lentiviral vector containing EGFP downstream of the human PGK promoter (a kind gift of Brown laboratory, ISMMS). Lentivirus was produced as described above. Cas9 expressing cells were then infected with pLenti-GFP-sgRNA.

### Cell proliferation assay

Cells were plated at a density of 5,000 cells/well with five replicates in a 96-well plate; four identical plates were prepared. Cell viability was assayed at 0, 2, 4 and 6 days after plating with alamarBlue Cell Viability Reagent (Thermo Fisher Scientific) and fluorescence at 585 nm was measured on a Spectra Max3 plate reader (Molecular Device) according to the manufacturer’s protocol at excitation of 555 nm. Cell viability at 2, 4 and 6 days were corrected for the ratio to control cells from the day 0 reading to account for plating unevenness.

### RNA-seq

Total RNAs from engineered NCI-H82 and NCI-H1963 cell lines were extracted using RNeasy kit (Qiagen). Poly-adenylated RNA was enriched from 1ug of RNA for each sample with the NEBNext^®^ PolyA mRNA Magnetic Isolation Module (NEB), incubated at 94°C for 15 min and double-strand cDNA was synthesized using SuperScript III reverse transcriptase (Thermo Fisher Scientific) and NEBNext^®^ Ultra™ II Directional RNA Second Strand Synthesis Module (NEB). Up to 10 ng of cDNA was used for the Illumina sequencing library construction using NEBNext^®^ Ultra™ DNA Library Prep Kit (NEB). Paired ends sequencing was performed on NextSeq 500 (Illumina) for 38 nucleotides from each end according to the manufacturer’s instructions.

### RNA-seq analyses

Sequencing reads were pseudoaligned to the human reference transcriptome GRCh38.95 from Ensembl and transcript abundance was estimated using kallisto (v0.45.0) (*48*). Transcript abundance was aggregated to gene level abundance using biomaRt annotation. DEseq2 (*49*) was used to identify differentially expressed genes between control NCI-H82 cells (LacZ-overexpressed NCI-H82 cells, LacZ-overexpressed + nontarget sgRNA1 NCI-H82 and LacZ-overexpressed + nontarget sgRNA2 NCI-H82) and L-Myc-overexpressed NCI-H82 cells (L-Myc-overexpressed NCI-H82 cells, L-Myc-overexpressed + nontarget sgRNA1 NCI-H82 and L-Myc-overexpressed + nontarget sgRNA2 NCI-H82). The same strategy was used for L-Myc-overexpressed NCI-H82 cells (L-Myc-overexpressed NCI-H82 cells, L-Myc-overexpressed + nontarget sgRNA1 NCI-H82 and L-Myc-overexpressed + nontarget sgRNA2 NCI-H82) and c-Myc replaced with L-Myc NCI-H82 cells (L-Myc-overexpressed NCI-H82 + sgRNA-MYC1 cells, L-Myc-overexpressed + sgRNA-MYC2 NCI-H82 and L-Myc-overexpressed + sgRNA-MYC3 NCI-H82), as well as for LacZ-overexpressed NCI-H1963 cells and c-Myc overexpressed NCI-H1963 cells.

Total differentially expressed genes were determined based on cutoffs of fold change> 2 and FDR<0.05. To identify potentially enriched functions of selected gene sets of interest, we compared these gene sets with the genes annotated by the same Gene Ontology (GO) terms curated in the Molecular Signature Database (MSigDB). Each of 5917 GO terms included in “C5” collection (version 7.0) was compared with query gene sets using Fisher’s exact test. The significance of the overlap was determined based on p-value adjusted for multiple comparisons (FDR<0.05). Any GO terms consisting of more than 2,000 genes were considered non-specific and removed from the analysis.

### Beta-galactosidase staining

Cells were stained with Senescence beta-galactosidase Staining Kit (Cell signaling #9860) according to manufacturer’s recommendations.

### Cell cycle analysis

Cells were harvested at day 2 after selection and fixed with 70 % ethanol overnight at 4°C, were washed with PBS, then, incubated in PBS containing 100 μg/ml RNase and 50 μg/ml propidium iodide at room-temperature for 1 hour. DNA content was analyzed by FACS Canto II (BD Bioscience), and quantitative analyses for the proportions of cells in cell cycle were performed using FlowJo software (BD Bioscience).

### Annexin-V staining

Cells were harvested at day 2 after selection and washed with PBS. Cells were stained with Annexin-V and propidium iodide using Alexa Fluor^®^ 488 annexin V/Dead Cell Apoptosis Kit (Thermo Fisher Scientific) according to manufacturers’ recommendations. Fluorescence was measured by FACS Canto II (BD Bioscience). Quantitative analyses for the cell viability proportions were performed using FlowJo software (BD Bioscience).

### Xenograft model

All animal procedures and studies were approved by the Mount Sinai Institutional Animal Care Use Committee (IACUC) (protocol number, IACUC-2018-0021). H1963 cells which overexpressed LacZ or cMYC (5 x 10^6^ cells) were injected with a 1:1 mixture of 50 μl cell suspension and 50 μl Matrigel (Corning) subcutaneously into both flank regions of 4-to 6-week-old male NOD-scid gamma mice (Jackson Laboratory). Tumor volume (length x width^2^ /2) was measured twice a week. When tumor size reached at 1000 mm^3^, mice were sacrificed and tumors were immersed in formalin for histological analysis.

### Immunohistochemistry

Xenograft tumor specimen formalin-fixed, paraffin-embedded tissue slides (5 μm thick) were deparaffinized and rehydrated. For antigen-retrieval slides were heated at 95°C in 10 mM citrate buffer (pH 6.0) for 30 minutes. The sections were incubated with 0.3% H2O2 in TBS for 15 minutes to block endogenous peroxidase activity and were incubated with 10% normal horse serum (Jackson ImmunoResearch) in TBS for 30 minutes to block non-specific staining. The sections were rinsed with TBS + 0.025% Triton X-100 and then incubated with anti-cMYC antibody (1:150; Abcam #ab32072) or anti-ASCL1 (1:100, BD Biosciences #556604) at 4°C overnight. This was followed by incubation with biotin-conjugated Horse anti-Mouse secondary antibody (Vector Laboratories) at room temperature for 1 hour. Then, the sections were incubated with the ABC reagent (Vector Laboratories, CA) and visualized with ImmPACT-DAB Peroxidase Substrate (Vector Laboratories). All slides were counterstained with hematoxylin before being mounting.

### Aurora Kinase Inhibition Assay

Cells were seeded in a 96-well plate at a density of 5,000 cells/well with five replicates. Cells were treated with Alisertib at concentrations ranging from 1 nM to 10 μM for 96h. Cell viability was measured using alamarBlue Cell Viability Reagent (Thermo Fisher Scientific) and fluorescence at 585nm was measured on a Spectra Max3 plate reader (Molecular Devices, CA) according to the manufacturer’s protocol at excitation of 555 nm.

## Acknowledgements

We thank Dawei Yang, Bhavana Shewale, Nicole Stokes, David Dominique-Sola, Christie Nguyen, Yifei Sun for helpful discussions. Aleksandra Wroblewska and Brian D. Brown for providing a lentiviral vector for cloning sgRNAs; NextSeq Sequencing Facility of the Department of Oncological Sciences at Icahn School of Medicine at Mount Sinai (ISMMS), Saboor Hekmaty, Gayatri Panda and Ravi Sachidanandam for sequencing assistance. The authors also thank the Flow Cytometry Core facility and the Biorepository and Pathology Core Facility at ISMMS. This work was supported in part through Tisch Cancer Institute at ISMMS and the computational resources and staff expertise provided by Scientific Computing at ISMMS. H. Watanabe is supported by 2017 ATS Foundation Unrestricted Grant, the American Lung Association of the Northeast, Department of Defense (W81XWH-19-1-0613) and NIH (R01CA240342). T. Sato is supported by the Japanese Respiratory Society the 6th Lilly Oncology Fellowship Program and the Uehara Memorial Foundation. R. Kong is supported by Shaanxi Provincial Natural Science Foundation, China (2017JM8046). C.A. Powell and H. Watanabe are supported by NIH (R01CA163772).

## Author Contribution

Conceptualization & Methodology, A.S.P. and H.W.; Data Collection: A.S.P., S.Y., R.K., T.S., A.S., L.B., M.F., K.E. and H.W.; Resources, M.B.B., C.A.P. and H.W.; Data Analysis, A.S.P., S.Y., G.N., M.B.B. and H.W.; Writing – Original Draft, A.S.P. and H.W.; Writing – Review & Editing, A.S.P., S.Y., G.N., C.A.P., M.B.B, J.Z. and H.W.; Supervision, C.A.P., J.Z. and H.W.; Project Administration & Funding Acquisition, H.W.

## Supplementary Figure Legends

**Figure S1:** Myc family status and signature in SCLC CCLE cell lines

A. Classification of 49 SCLC cell lines using copy number and expression of *MYC, MYCL* and *MYCN* from CCLE

B. Scatter plot showing c-Myc expression and L-Myc expression for 49 SCLC cell lines. Cell lines selected to generate signature are highlighted L-Myc in pink and c-Myc in blue.

C. Heatmap showing 457 differentially expressed (t-test p-value < 0.01 and fold change > 1.5) between four c-Myc expressing CCLE SCLC cell lines (H1341, H2081, H211, H524, SCLC21H) and eight L-Myc expressing CCLE SCLC cell lines (HCC33, H1092, H1184, H1836, H1963, H2029, H209). 147 genes upregulated in c-Myc (condition shown in blue) and 310 genes upregulated in L-Myc (condition shown in pink).

**Figure S2**: ATAC-seq inferred c-Myc accessibility is comparable to c-Myc ChIP-seq detected c-Myc bound regions and classifies by Myc status

A. Heatmap comparing Myc (E-box) accessibility inferred from ATAC-seq with c-Myc bound sites with E-box motif detected from ChIP-seq (Lin et al., 2012) in NCI-H2171. ATAC-seq/ChIP-seq signal intensity is shown by color shading.

B. Genome browser view tracks of ATAC-seq signals and c-Myc ChIP-seq at the *HES1* locus.

C. Heatmap showing unsupervised clustering of degree of correlation for pairwise comparisons of identified open chromatin regions with *MYC* motif in 3 L-Myc classified cell lines H209, H1963, CORL88 (condition shown in pink) and 3 c-Myc classified cell lines H524, H82, H2171 (condition shown in blue)

**Figure S3**: c-Myc and L-Myc expression across SCLC molecular subtypes

A-D Genome browser view tracks of ATAC-seq signals in c-Myc cell lines shown in blue (NCI-H82, NCI-H524, NCI-H2171) and L-Myc cell lines peaks shown in pink (NCI-H209, NCI-H1963, CORL88) at the (A) *NeuroD1,* (B) *ASCL1,* (C) *POU2F3,* (D) YAP1.

E. Immunoblot showing protein expression of NeuroD1 and ASCL1 in c-Myc and L-Myc classified cell lines with Vinculin as loading control.

F. Violin plot showing mRNA expression of c-Myc across four molecular subtypes in CCLE cell lines as classified in Rudin et al., 2019

G. Violin plot showing mRNA expression of L-Myc across four molecular subtypes in CCLE cell lines as classified in Rudin et al., 2019

**Figure S4**: L-Myc induces neuronal state not defined by ASCL-1

A. Hematoxylin-eosin staining in NCI-H82 LacZ-overexpressed with nontarget gRNA (left panel), NCI-H82 L-Myc-overexpressed with nontarget gRNA (middle panel) or NCI-H82 L-Myc-overexpressed with gRNA targeting *MYC* (Right panel) xenograft. Original images, x20.

B. Volcano plot showing 525 differentially expressed genes (235 upregulated in purple and 290 downregulated in blue) between control NCI-H82 cells (LacZ-overexpressed cells and LacZ-overexpressed cells with nontarget gRNA) and engineered NCI-H82 cells (LMYC-overexpressed cells and LMYC-overexpressed cells with nontarget gRNA).

C. Gene Ontology analysis for differentially upregulated (top panel) and downregulated (bottom panel) genes from Figure S4B.

D. Volcano plot showing 214 differentially expressed genes (170 upregulated in pink and 44 downregulated in purple) between L-Myc-overexpressed NCI-H82 cells (L-Myc-overexpressed cells and L-Myc-overexpressed cells with nontarget sgRNA) and Myc-replaced NCI-H82 cells (L-Myc-overexpressed with three sgRNA targeting MYC).

E. Gene Ontology analysis for differentially upregulated (top panel) and downregulated (bottom panel) genes from Figure S4D.

**Figure S5:** Aurora Kinase A inhibition shows differential sensitivity between c-Myc and L-Myc expressing cells

A. Heatmap showing dose-dependent anti-proliferative activity of Alisertib as screened across a panel of 3 c-Myc (shown in blue) and 3 L-Myc cell lines (shown in pink) at 96 hours. Cell viability (Alamar Blue) was calculated relative to the control. Data are means from at least *n* = 3 biological replicates.

B. Dose-response curves of H82-replacement cells treated with Alisertib for 96 hours. Viability was assessed with the Alamar Blue assay and calculated relative to the control. Data are means from at least *n* = 3 biological replicates.

C. Dose-response curves of H1963–genetically engineeredt cells treated with Alisertib for 96 hours. Viability was assessed with the AlamarBlue assay and calculated relative to the control. Data are means from at least *n* = 3 biological replicates.

**Figure S6**: c-Myc induced growth-suppression is not a consequence of altered cell cycle dynamics

A. Beta-galactosidase staining for c-Myc-overexpressed NCI-H1963 cells (bottom panel) or control LacZ-overexpressed NCI-H1963 (top panel) or at day 2.

B. Propidium iodide staining followed by flow cytometry showing cell cycle distribution in c-Myc overexpressed NCI-H1963 cells and control LacZ-overexpressed NCI-H1963 LacZ-overexpressed control cells at day 2.

C. Hematoxylin-eosin staining in LacZ-overexpressed (left) or c-Myc-overexpressed (right) NCI-H1963 xenograft. Original Images, ×5 (top) and x20 (bottom).

