## Supplementary Figures and Tables for "Classic oncogene family Myc defines unappreciated distinct lineage states of small cell lung cancer"

### Supplementary Figure 1

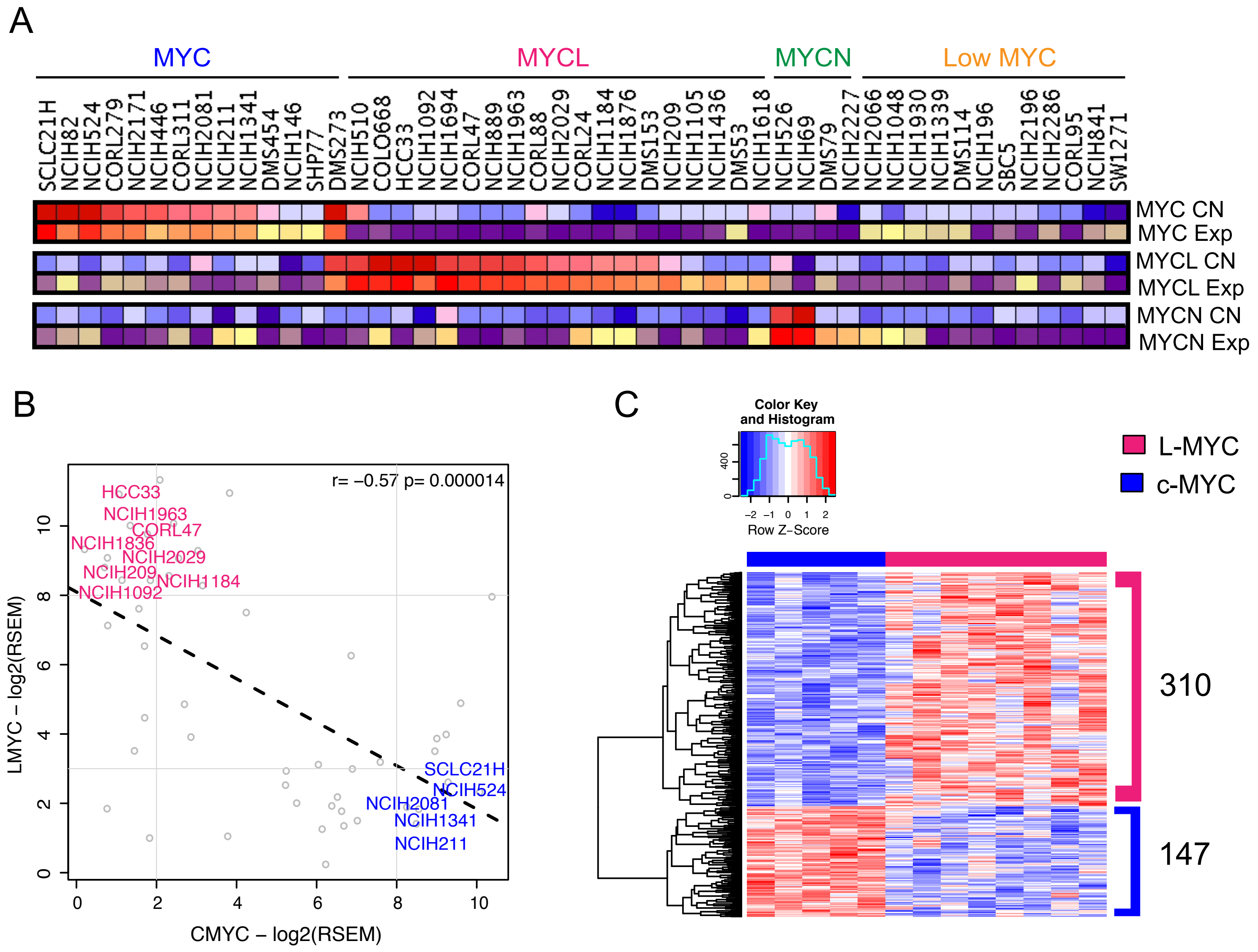

### Supplementary Figure 2

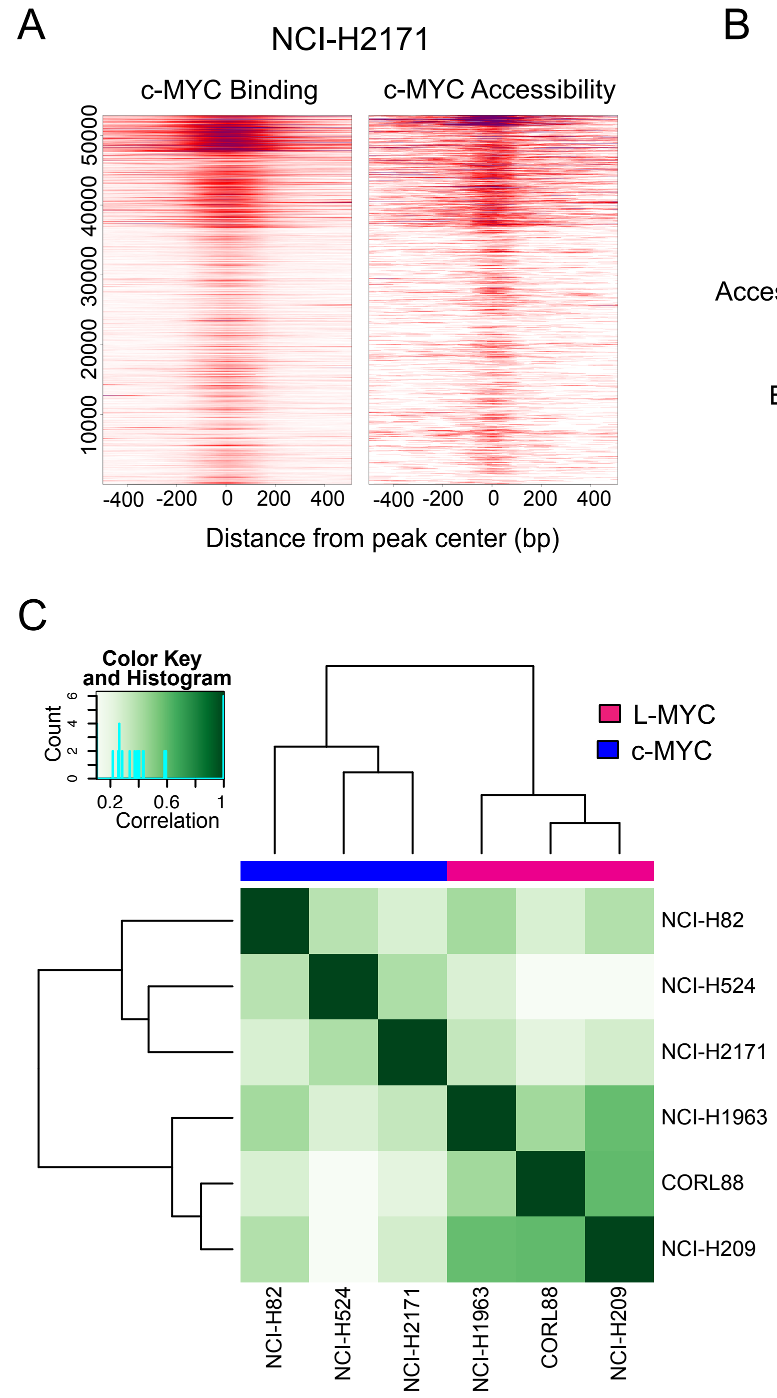

### Supplementary Figure 3

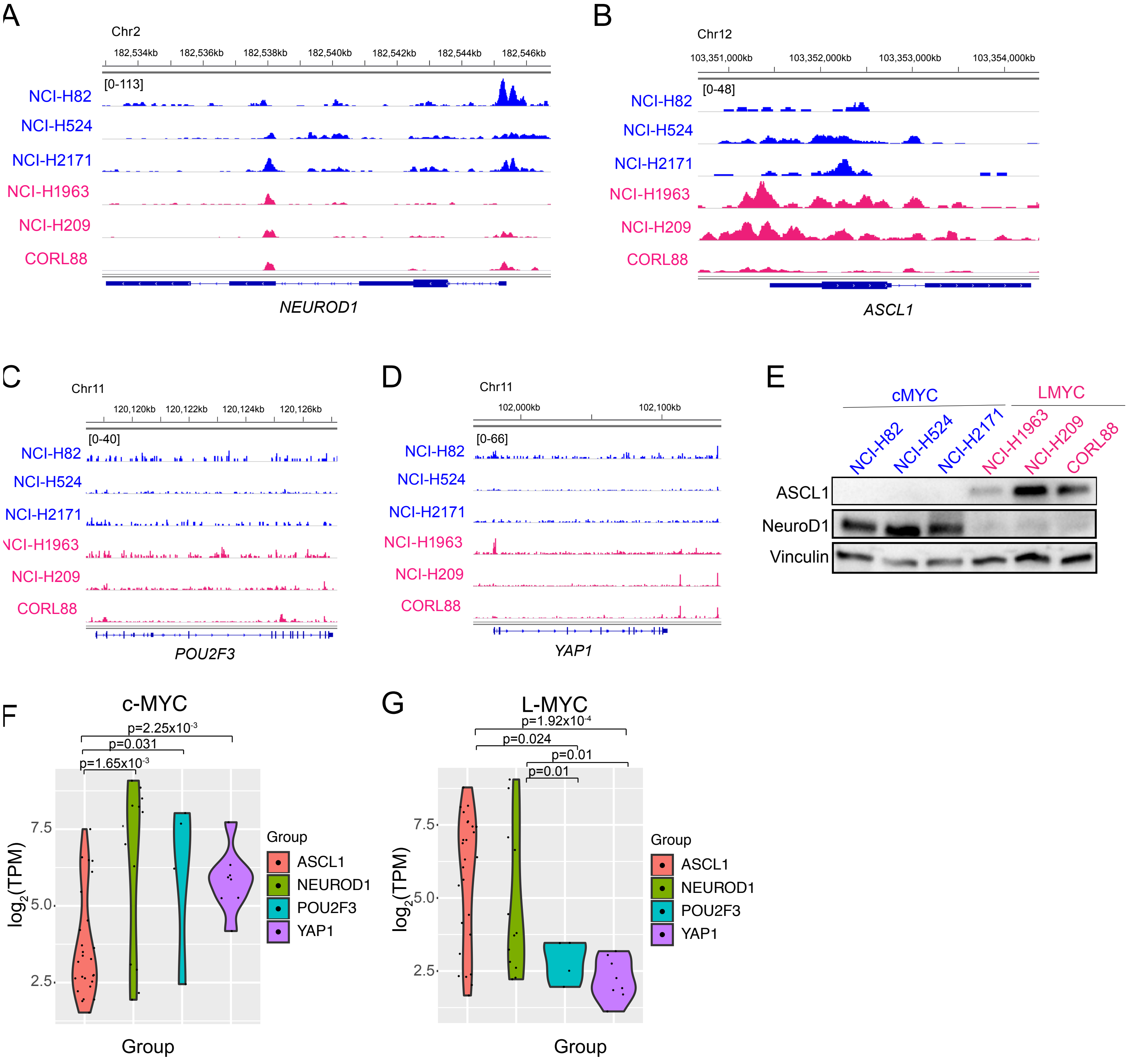

Supplementary Figure 4

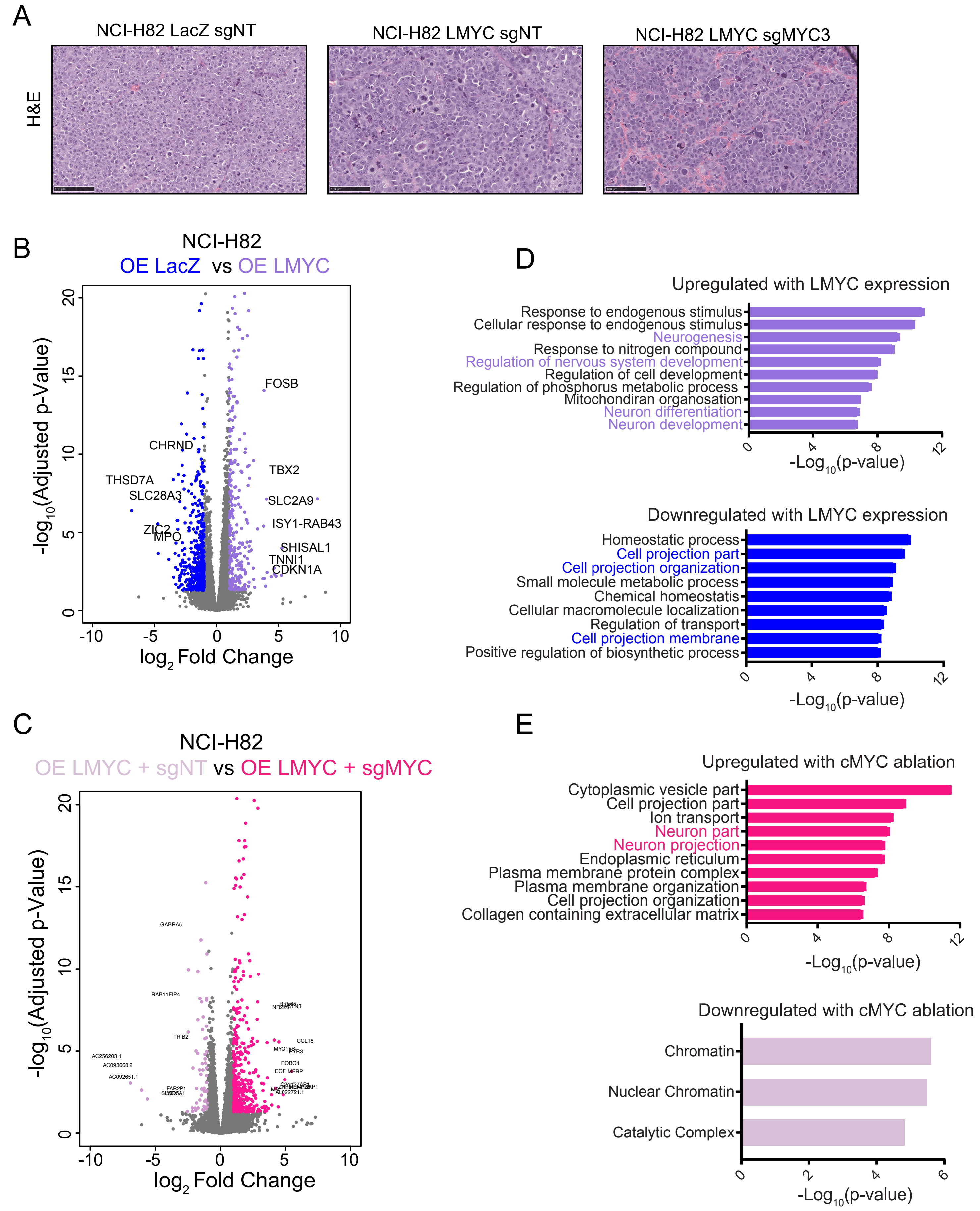

### Supplementary Figure 5

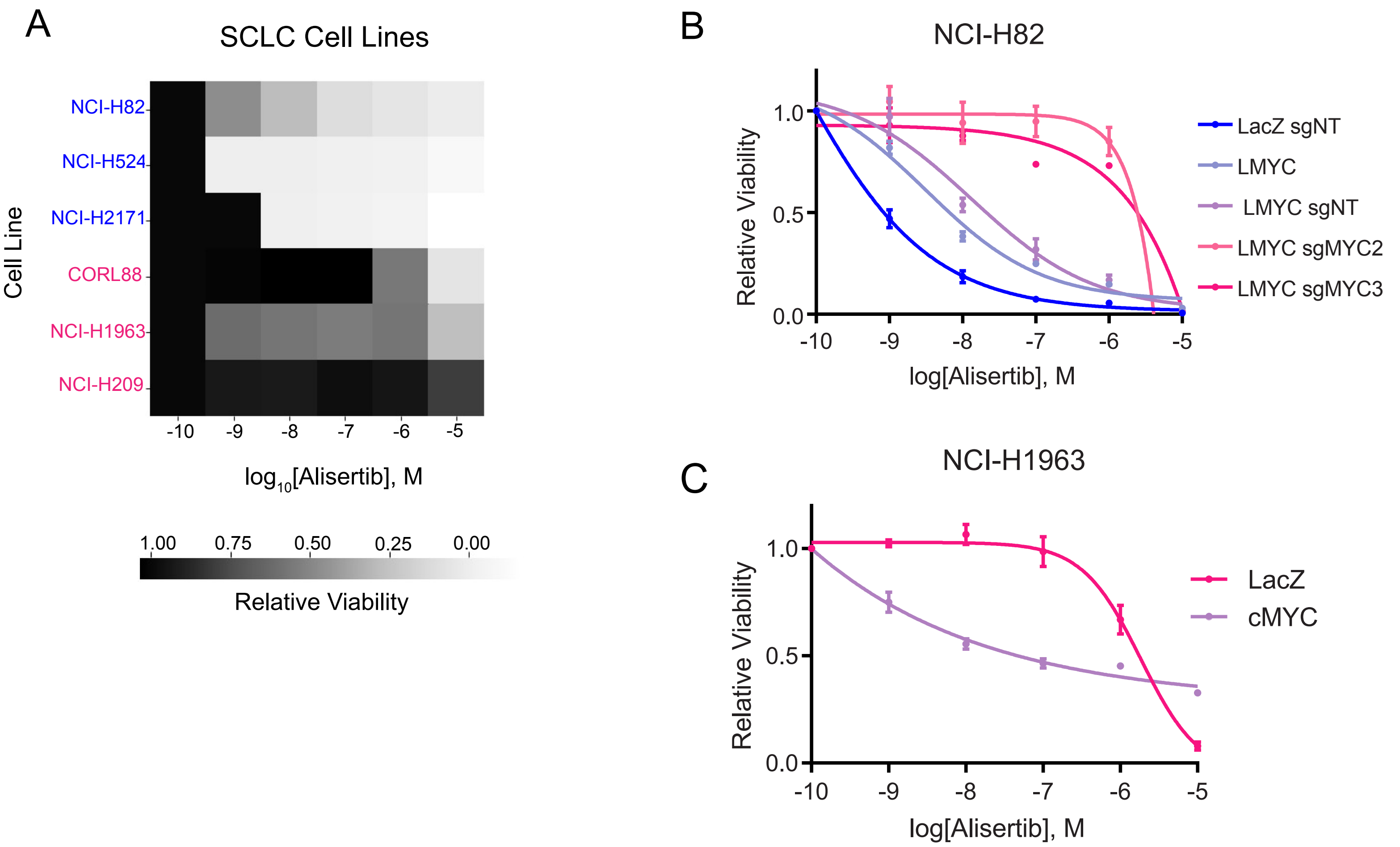

#### Table S1: Key drivers for LMYC Network

s

| Gene | Target | Overlap | OR | p-Value |
|---|---|---|---|---|
| *NFYB* | 3 | 3 | Inf | 7.86E-06 |
| *TM9SF2* | 3 | 3 | Inf | 7.86E-06 |
| *PABPC4* | 3 | 3 | Inf | 7.86E-06 |
| *ABDH13* | 5 | 3 | 74.46 | 7.63E-05 |
| *MCF2L* | 13 | 4 | 22.24 | 9.58E-05 |
| *CBFA2T2* | 14 | 4 | 20.02 | 1.32E-04 |
| *CACNA1A* | 26 | 5 | 11.98 | 1.41E-04 |
| *SMAD2* | 18 | 4 | 14.3 | 3.79E-04 |
| *FAM127C* | 2 | 2 | Inf | 3.98E-04 |
| *EPG5* | 2 | 2 | Inf | 3.98E-04 |
| *RAP2A* | 8 | 3 | 29.83 | 4.09E-04 |
| *ADARB2-AS1* | 10 | 3 | 21.33 | 8.50E-04 |
| *CAMK1D* | 10 | 3 | 21.33 | 8.50E-04 |

**Table S2: Key drivers for cMYC Network (subnetwork c1- c3)**

| Gene | Target | Overlap | OR | p-Value |
|---|---|---|---|---|
| *PLEKH02* | 17 | 3 | 31.97 | 2.05E-04 |
| *RPL18* | 19 | 3 | 27.98 | 2.89E-04 |
| *ANXA1* | 21 | 3 | 24.85 | 3.93E-04 |
| *RPS3* | 21 | 3 | 24.85 | 3.93E-04 |
| *FAM114A1* | 5 | 2 | 97.27 | 4.73E-04 |
| *NPTX2* | 5 | 2 | 97.27 | 4.73E-04 |
| *STX4* | 5 | 2 | 97.27 | 4.73E-04 |
| *MMP14* | 5 | 2 | 97.27 | 4.73E-04 |
| *AHNAK* | 24 | 3 | 21.32 | 5.89E-04 |
| *CASP4* | 6 | 2 | 73.17 | 7.06E-04 |
| *DHRS3* | 6 | 2 | 73.17 | 7.06E-04 |
| *LHFPL4* | 6 | 2 | 73.17 | 7.06E-04 |
| *NAB2* | 7 | 2 | 58.5 | 9.84E-04 |
| *WDR43* | 7 | 2 | 58.5 | 9.84E-04 |
| *HSP90Ab1* | 7 | 2 | 58.5 | 9.84E-04 |

**Table S3:** sgRNA sequences for the CRISPR/Cas9 system

| | sgRNA (5'-3') |
|---|---|
| MYC sgRNA1 | AGAGTGCATCGACCCCTCGG |
| MYC sgRNA2 | CTGCGGGGAGGACTCCGTCG |
| MYC sgRNA3 | CTTCGGGGAGACAACGACGG |
| Nontarget sgRNA 1 | ACGGAGGCTAAGCGTCGCAA |
| Nontarget sgRNA 2 | CGCTTCCGCGGCCCGTTCAA |

**Table S4: Primary Tumor and CCLE Cell Line Subtype Classification**

| Name | Source | Tumor or Cell line | Subtype Classification (This Paper) | Subtype Classification (Rudin et al.2019) |
|---|---|---|---|---|
| SRR1797249 | Jiang et al. 2016 | Tumor | ASCL1 | NA |
| SRR1797247 | Jiang et al. 2016 | Tumor | ASCL1 | NA |
| SRR1797267 | Jiang et al. 2016 | Tumor | ASCL1 | NA |
| S01297T | George et al. 2015 | Tumor | ASCL1 | ASCL1 |
| SRR1797271 | Jiang et al. 2016 | Tumor | ASCL1 | NA |
| SRR1797294 | Jiang et al. 2016 | Tumor | ASCL1 | NA |
| SRR1797269 | Jiang et al. 2016 | Tumor | ASCL1 | NA |
| SRR1797231 | Jiang et al. 2016 | Tumor | ASCL1 | NA |
| SRR1797260 | Jiang et al. 2016 | Tumor | ASCL1 | NA |
| CORL279 | CCLE | Cell line | ASCL1 | NEUROD1 |
| SRR1797286 | Jiang et al. 2016 | Tumor | ASCL1 | NA |
| S01242 | George et al. 2015 | Tumor | ASCL1 | NA |
| S00832 | George et al. 2015 | Tumor | ASCL1 | ASCL1 |
| S02360 | George et al. 2015 | Tumor | ASCL1 | ASCL1 |
| S02065 | George et al. 2015 | Tumor | ASCL1 | ASCL1 |
| SRR1797232 | Jiang et al. 2016 | Tumor | ASCL1 | NA |
| NCIH2066 | CCLE | Cell line | ASCL1 | NEUROD1 |
| SRR1797265 | Jiang et al. 2016 | Tumor | ASCL1 | NA |
| SRR1797270 | Jiang et al. 2016 | Tumor | ASCL1 | NA |
| S02378T | George et al. 2015 | Tumor | ASCL1 | ASCL1 |
| SRR1797236 | Jiang et al. 2016 | Tumor | ASCL1 | NA |
| SRR1797261 | Jiang et al. 2016 | Tumor | ASCL1 | NA |
| NCIH2227 | CCLE | Cell line | ASCL1 | NEUROD1 |
| S02243 | George et al. 2015 | Tumor | ASCL1 | ASCL1 |
| S02241 | George et al. 2015 | Tumor | ASCL1 | ASCL1 |
| SRR1797228 | Jiang et al. 2016 | Tumor | ASCL1 | NA |
| SRR1797292 | Jiang et al. 2016 | Tumor | ASCL1 | NA |
| S02291 | George et al. 2015 | Tumor | ASCL1 | ASCL1 |
| SRR1797273 | Jiang et al. 2016 | Tumor | ASCL1 | NA |
| SRR1797303 | Jiang et al. 2016 | Tumor | ASCL1 | NA |
| SRR1797225 | Jiang et al. 2016 | Tumor | ASCL1 | NA |
| NCIH1876 | CCLE | Cell line | ASCL1 | ASCL1 |
| SRR1797281 | Jiang et al. 2016 | Tumor | ASCL1 | NA |
| SRR1797288 | Jiang et al. 2016 | Tumor | ASCL1 | NA |
| S02284 | George et al. 2015 | Tumor | ASCL1 | ASCL1 |
| NCIH1618 | CCLE | Cell line | ASCL1 | ASCL1 |
| SRR1797241 | Jiang et al. 2016 | Tumor | ASCL1 | NA |
| S02297 | George et al. 2015 | Tumor | ASCL1 | ASCL1 |
| S02290 | George et al. 2015 | Tumor | ASCL1 | ASCL1 |
| S02376T | George et al. 2015 | Tumor | ASCL1 | ASCL1 |

| | | | | |
|---|---|---|---|---|
| SRR1797274 | Jiang et al. 2016 | Tumor | ASCL1 | NA |
| NCIH889 | CCLE | Cell line | ASCL1 | ASCL1 |
| SRR1797300 | Jiang et al. 2016 | Tumor | ASCL1 | NA |
| S00831 | George et al. 2015 | Tumor | ASCL1 | ASCL1 |
| S00825 | George et al. 2015 | Tumor | ASCL1 | ASCL1 |
| SRR1797277 | Jiang et al. 2016 | Tumor | ASCL1 | NA |
| SRR1797239 | Jiang et al. 2016 | Tumor | ASCL1 | NA |
| S01864 | George et al. 2015 | Tumor | ASCL1 | ASCL1 |
| SRR1797295 | Jiang et al. 2016 | Tumor | ASCL1 | NA |
| SRR1797283 | Jiang et al. 2016 | Tumor | ASCL1 | NA |
| DMS454 | CCLE | Cell line | ASCL1 | ASCL1 |
| S01366 | George et al. 2015 | Tumor | ASCL1 | ASCL1 |
| SRR1797242 | Jiang et al. 2016 | Tumor | ASCL1 | NA |
| SRR1797276 | Jiang et al. 2016 | Tumor | ASCL1 | NA |
| S00035T | George et al. 2015 | Tumor | ASCL1 | ASCL1 |
| S02255 | George et al. 2015 | Tumor | ASCL1 | NA |
| S02249 | George et al. 2015 | Tumor | ASCL1 | ASCL1 |
| SRR1797255 | Jiang et al. 2016 | Tumor | ASCL1 | NA |
| NCIH510 | CCLE | Cell line | ASCL1 | ASCL1 |
| S02285 | George et al. 2015 | Tumor | ASCL1 | ASCL1 |
| S02295 | George et al. 2015 | Tumor | ASCL1 | ASCL1 |
| SRR1797251 | Jiang et al. 2016 | Tumor | ASCL1 | NA |
| SRR1797289 | Jiang et al. 2016 | Tumor | ASCL1 | NA |
| SRR1797279 | Jiang et al. 2016 | Tumor | ASCL1 | NA |
| S02287 | George et al. 2015 | Tumor | ASCL1 | ASCL1 |
| CORL47 | CCLE | Cell line | ASCL1 | ASCL1 |
| NCIH1930 | CCLE | Cell line | ASCL1 | ASCL1 |
| SHP77 | CCLE | Cell line | ASCL1 | ASCL1 |
| CORL95 | CCLE | Cell line | ASCL1 | ASCL1 |
| NCIH2029 | CCLE | Cell line | ASCL1 | ASCL1 |
| NCIH146 | CCLE | Cell line | ASCL1 | ASCL1 |
| NCIH1105 | CCLE | Cell line | ASCL1 | ASCL1 |
| NCIH1963 | CCLE | Cell line | ASCL1 | ASCL1 |
| S02299 | George et al. 2015 | Tumor | ASCL1 | NA |
| NCIH1436 | CCLE | Cell line | ASCL1 | ASCL1 |
| NCIH2196 | CCLE | Cell line | ASCL1 | ASCL1 |
| S02328 | George et al. 2015 | Tumor | ASCL1 | ASCL1 |
| S02194 | George et al. 2015 | Tumor | ASCL1 | ASCL1 |
| SRR1797285 | Jiang et al. 2016 | Tumor | ASCL1 | NA |
| SRR1797264 | Jiang et al. 2016 | Tumor | ASCL1 | NA |
| S00213 | George et al. 2015 | Tumor | ASCL1 | ASCL1 |
| SRR1797234 | Jiang et al. 2016 | Tumor | ASCL1 | NA |
| SRR1797227 | Jiang et al. 2016 | Tumor | ASCL1 | NA |
| SRR1797301 | Jiang et al. 2016 | Tumor | ASCL1 | NA |
| SRR1797268 | Jiang et al. 2016 | Tumor | ASCL1 | NA |

| | | | | |
|---|---|---|---|---|
| SRR1797244 | Jiang et al. 2016 | Tumor | ASCL1 | NA |
| SRR1797298 | Jiang et al. 2016 | Tumor | ASCL1 | NA |
| SRR1797275 | Jiang et al. 2016 | Tumor | ASCL1 | NA |
| S02248 | George et al. 2015 | Tumor | ASCL1 | ASCL1 |
| SRR1797259 | Jiang et al. 2016 | Tumor | ASCL1 | NA |
| S01578 | George et al. 2015 | Tumor | ASCL1 | ASCL1 |
| SRR1797291 | Jiang et al. 2016 | Tumor | ASCL1 | NA |
| S00022 | George et al. 2015 | Tumor | ASCL1 | ASCL1 |
| SRR1797229 | Jiang et al. 2016 | Tumor | ASCL1 | NA |
| NCIH1092 | CCLE | Cell line | ASCL1 | ASCL1 |
| NCIH2081 | CCLE | Cell line | ASCL1 | ASCL1 |
| SRR1797257 | Jiang et al. 2016 | Tumor | ASCL1 | NA |
| S02244 | George et al. 2015 | Tumor | ASCL1 | ASCL1 |
| SRR1797256 | Jiang et al. 2016 | Tumor | ASCL1 | NA |
| SRR1797280 | Jiang et al. 2016 | Tumor | ASCL1 | NA |
| S02397 | George et al. 2015 | Tumor | ASCL1 | ASCL1 |
| SRR1797235 | Jiang et al. 2016 | Tumor | ASCL1 | NA |
| S02289 | George et al. 2015 | Tumor | ASCL1 | ASCL1 |
| SRR1797238 | Jiang et al. 2016 | Tumor | ASCL1 | NA |
| S02242 | George et al. 2015 | Tumor | ASCL1 | ASCL1 |
| S02322 | George et al. 2015 | Tumor | ASCL1 | ASCL1 |
| DMS53 | CCLE | Cell line | ASCL1 | ASCL1 |
| SRR1797272 | Jiang et al. 2016 | Tumor | ASCL1 | NA |
| S00838 | George et al. 2015 | Tumor | ASCL1 | ASCL1 |
| SRR1797233 | Jiang et al. 2016 | Tumor | ASCL1 | NA |
| SRR1797246 | Jiang et al. 2016 | Tumor | ASCL1 | NA |
| S02093 | George et al. 2015 | Tumor | ASCL1 | NA |
| S02120 | George et al. 2015 | Tumor | ASCL1 | ASCL1 |
| SRR1797284 | Jiang et al. 2016 | Tumor | ASCL1 | NA |
| SRR1797245 | Jiang et al. 2016 | Tumor | ASCL1 | NA |
| NCIH1184 | CCLE | Cell line | ASCL1 | ASCL1 |
| SRR1797237 | Jiang et al. 2016 | Tumor | ASCL1 | NA |
| SRR1797282 | Jiang et al. 2016 | Tumor | ASCL1 | NA |
| S02382T | George et al. 2015 | Tumor | ASCL1 | ASCL1 |
| SRR1797299 | Jiang et al. 2016 | Tumor | ASCL1 | NA |
| SRR1797263 | Jiang et al. 2016 | Tumor | ASCL1 | NA |
| SRR1797240 | Jiang et al. 2016 | Tumor | ASCL1 | NA |
| S01524 | George et al. 2015 | Tumor | ASCL1 | ASCL1 |
| SRR1797250 | Jiang et al. 2016 | Tumor | ASCL1 | NA |
| SRR1797293 | Jiang et al. 2016 | Tumor | ASCL1 | NA |
| DMS79 | CCLE | Cell line | ASCL1 | ASCL1 |
| NCIH1836 | CCLE | Cell line | ASCL1 | ASCL1 |
| CORL88 | CCLE | Cell line | ASCL1 | ASCL1 |
| S01861T | George et al. 2015 | Tumor | ASCL1 | ASCL1 |
| S01512 | George et al. 2015 | Tumor | ASCL1 | ASCL1 |

| | | | | |
|---|---|---|---|---|
| SRR1797243 | Jiang et al. 2016 | Tumor | ASCL1 | NA |
| COLO668 | CCLE | Cell line | ASCL1 | ASCL1 |
| NCIH209 | CCLE | Cell line | ASCL1 | ASCL1 |
| SRR1797262 | Jiang et al. 2016 | Tumor | ASCL1 | NA |
| NCIH69 | CCLE | Cell line | ASCL1 | ASCL1 |
| DMS153 | CCLE | Cell line | ASCL1 | ASCL1 |
| S02375T | George et al. 2015 | Tumor | NEUROD1 | POU2F3 |
| SRR1797302 | Jiang et al. 2016 | Tumor | NEUROD1 | NA |
| SRR1797253 | Jiang et al. 2016 | Tumor | NEUROD1 | NA |
| SRR1797266 | Jiang et al. 2016 | Tumor | NEUROD1 | NA |
| S01873T | George et al. 2015 | Tumor | NEUROD1 | NEUROD1 |
| NCIH446 | CCLE | Cell line | NEUROD1 | NEUROD1 |
| NCIH524 | CCLE | Cell line | NEUROD1 | NEUROD1 |
| DMS273 | CCLE | Cell line | NEUROD1 | NEUROD1 |
| CORL24 | CCLE | Cell line | NEUROD1 | NEUROD1 |
| S02139 | George et al. 2015 | Tumor | NEUROD1 | NEUROD1 |
| NCIH1694 | CCLE | Cell line | NEUROD1 | NEUROD1 |
| SRR1797278 | Jiang et al. 2016 | Tumor | NEUROD1 | NA |
| SCLC21H | CCLE | Cell line | NEUROD1 | NEUROD1 |
| S02163 | George et al. 2015 | Tumor | NEUROD1 | NEUROD1 |
| HCC33 | CCLE | Cell line | NEUROD1 | NEUROD1 |
| NCIH82 | CCLE | Cell line | NEUROD1 | NEUROD1 |
| NCIH2171 | CCLE | Cell line | NEUROD1 | NEUROD1 |
| S02294 | George et al. 2015 | Tumor | NEUROD1 | NEUROD1 |
| S02293 | George et al. 2015 | Tumor | NEUROD1 | NEUROD1 |
| S00829 | George et al. 2015 | Tumor | POU2F3 | POU2F3 |
| S02288 | George et al. 2015 | Tumor | POU2F3 | POU2F3 |
| SRR1797254 | Jiang et al. 2016 | Tumor | POU2F3 | NA |
| SRR1797297 | Jiang et al. 2016 | Tumor | POU2F3 | NA |
| NCIH1048 | CCLE | Cell line | POU2F3 | POU2F3 |
| S02234 | George et al. 2015 | Tumor | POU2F3 | NA |
| SRR1797230 | Jiang et al. 2016 | Tumor | POU2F3 | NA |
| S01542 | George et al. 2015 | Tumor | POU2F3 | NA |
| S02209 | George et al. 2015 | Tumor | POU2F3 | POU2F3 |
| S02286 | George et al. 2015 | Tumor | POU2F3 | POU2F3 |
| S02296 | George et al. 2015 | Tumor | POU2F3 | POU2F3 |
| S02256 | George et al. 2015 | Tumor | POU2F3 | POU2F3 |
| SRR1797258 | Jiang et al. 2016 | Tumor | POU2F3 | NA |
| SRR1797248 | Jiang et al. 2016 | Tumor | POU2F3 | NA |
| NCIH211 | CCLE | Cell line | POU2F3 | POU2F3 |
| CORL311 | CCLE | Cell line | POU2F3 | POU2F3 |
| NCIH526 | CCLE | Cell line | POU2F3 | POU2F3 |
| SRR1797296 | Jiang et al. 2016 | Tumor | YAP1 | NA |
| S02298 | George et al. 2015 | Tumor | YAP1 | YAP1 |
| S02246 | George et al. 2015 | Tumor | YAP1 | YAP1 |

| | | | | |
|---|---|---|---|---|
| SRR1797290 | Jiang et al. 2016 | Tumor | YAP1 | NA |
| NCIH1341 | CCLE | Cell line | YAP1 | YAP1 |
| SW1271 | CCLE | Cell line | YAP1 | YAP1 |
| NCIH1339 | CCLE | Cell line | YAP1 | YAP1 |
| NCIH841 | CCLE | Cell line | YAP1 | YAP1 |
| SRR1797287 | Jiang et al. 2016 | Tumor | YAP1 | NA |
| DMS114 | CCLE | Cell line | YAP1 | YAP1 |
| NCIH196 | CCLE | Cell line | YAP1 | YAP1 |
| SBC5 | CCLE | Cell line | YAP1 | YAP1 |
| NCIH2286 | CCLE | Cell line | YAP1 | YAP1 |
